# Nutrient regulation of the islet epigenome controls adaptive insulin secretion

**DOI:** 10.1101/742403

**Authors:** Matthew Wortham, Fenfen Liu, Johanna Y. Fleischman, Martina Wallace, Francesca Mulas, Nicholas K. Vinckier, Austin R. Harrington, Benjamin R. Cross, Joshua Chiou, Nisha A. Patel, Yinghui Sui, Ulupi S. Jhala, Orian S. Shirihai, Mark O. Huising, Kyle J. Gaulton, Christian M. Metallo, Maike Sander

## Abstract

Adaptation of the islet β-cell insulin secretory response to changing insulin demand is critical for blood glucose homeostasis, yet the mechanisms underlying this adaptation are unknown. Here, we show that nutrient-stimulated histone acetylation plays a key role in adapting insulin secretion through regulation of genes involved in β-cell nutrient sensing and metabolism. Nutrient regulation of the epigenome occurs at sites occupied by the chromatin-modifying enzyme Lsd1 in islets. We demonstrate that β-cell-specific deletion of *Lsd1* leads to insulin hypersecretion, aberrant expression of nutrient response genes, and histone hyperacetylation. Islets from mice adapted to chronically increased insulin demand exhibited similar epigenetic and transcriptional changes. Moreover, genetic variants associated with fasting glucose and type 2 diabetes are enriched at LSD1-bound sites in human islets, suggesting that interpretation of nutrient signals is genetically determined. Our findings reveal that adaptive insulin secretion involves Lsd1-mediated coupling of nutrient state to regulation of the islet epigenome.

## Main

The ability to regulate nutrient metabolism in response to feeding and fasting is necessary for metabolic homeostasis. Nutrient utilization is acutely regulated by hormones and metabolites that change in response to feeding state^1^. If an energy state persists, adaptive control mechanisms increasingly influence nutrient metabolism. This is exemplified by adaptation of liver metabolism to fasting, wherein a network of nutrient- and hormone-responsive transcriptional complexes act in tandem with epigenetic changes to drive metabolic rewiring coupled to changes in fuel availability^2–6^. Understanding how feeding and fasting signals are interpreted in metabolic tissues is critical to developing a comprehensive understanding of energy homeostasis.

Insulin produced by pancreatic β-cells is the key stimulus for carbohydrate metabolism, and therefore it is critical that insulin secretion is adjusted commensurate with changes in energy state^1^. The insulin secretory response to feeding is primarily controlled by glucose and potentiated by intra-islet glucagon^7,8^ and incretin hormones such as Glp-1 through stimulation of cAMP production^9–11^. The postprandial increase of serum glucose accelerates glucose metabolism within the β-cell to initiate a signalling cascade involving ATP-stimulated plasma membrane depolarization, Ca^2+^ influx, and insulin vesicle exocytosis^12^. Although much is known regarding how nutritional and hormonal signals acutely regulate the insulin secretory response^9^, it is unclear how feeding and fasting evoke sustained functional changes as β-cells adapt to these nutrient states.

It has long been recognized that the insulin secretory response is attenuated by prolonged fasting^13–15^ and sensitized in response to nutrient overload during adaptation to obesity^16,17^. During fasting, β-cells rewire their metabolism to increase fatty acid oxidation at the expense of glucose metabolism, thereby reducing the ability of glucose to stimulate insulin secretion^18,19^. This metabolic switch is in part transcriptionally driven through upregulation of Pparα^20^. In the fed state, key nutritional signals that promote insulin secretion such as glucose and Glp-1 also trigger changes in β-cell gene expression^21,22^, raising the possibility that these feeding-induced transcriptional changes contribute to adaptive enhancement of the insulin secretory response. Gene regulation in response to glucose has also been associated with changes in histone acetylation^23,24^, suggesting that chromatin modifications could contribute to nutrient-regulated transcription in β-cells. It is unknown whether feeding is coupled to transcriptional or epigenomic regulation in β-cells to mediate functional changes that endure over the duration of the fed state.

To investigate adaptation of pancreatic islets to changes in nutrient state, we employed a time-restricted feeding paradigm^25^ to reinforce natural rhythms of food intake in mice. Food availability was restricted to the 12-hour dark phase for six days, which did not result in weight loss (**Extended Data Fig. 1a, b**). Notably, time-restricted feeding of normal chow has previously been shown to not affect glucose tolerance^25^. On the final day of entrainment after 12 hours of fasting, food was provided to one group for 4 hours and withheld from the fasted group, which resulted in differences in blood glucose levels (**Extended Data Fig. 1c**). Glucose stimulated insulin secretion (GSIS) by isolated islets was higher in the fed than in the fasted state (**Extended Data Fig. 1d**), suggesting that even short-term changes in nutrient state trigger adaptive changes in the insulin secretory response. To assess transcriptional changes associated with adaptation to feeding, we analysed islets from fasted and fed mice by mRNA-seq (**Fig. 1a**, **Supplementary Table 1**), revealing 1,186 differentially expressed genes (*P* < 0.01; **Extended Data Fig. 2a**, **Supplementary Table 2a**). Functional annotation of feeding-responsive genes identified regulation of metabolic (carbohydrate, lipid, and amino acid metabolism) and nutrient-sensing (MAPK, mTOR, and FoxO) pathways (**Fig. 1b, c**) known to adapt the insulin secretory response to energy state^17–19,26–30^.

**Figure 1.**
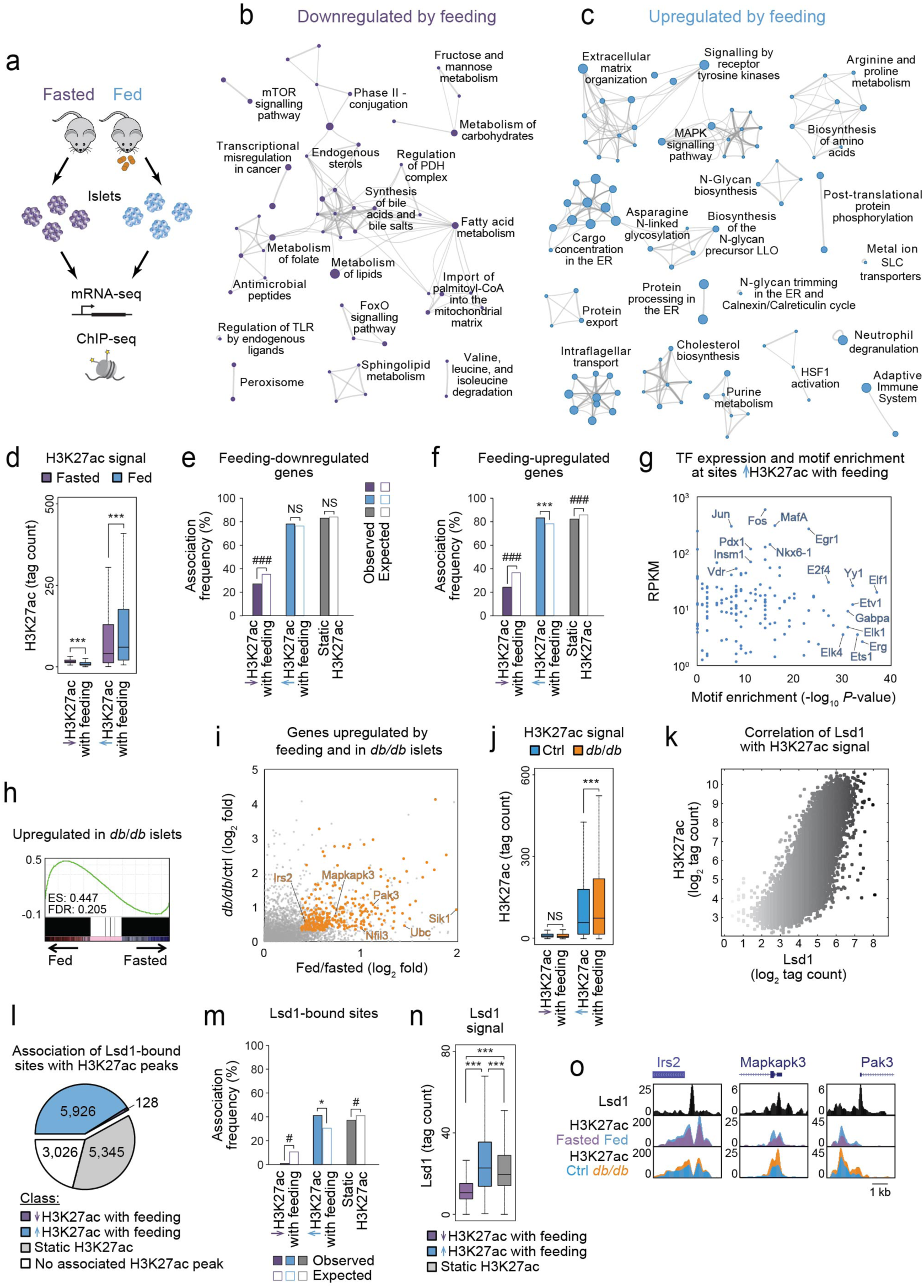
Nutrient state regulates histone acetylation and transcription in pancreatic islets. **(a)** Schematic of experiments performed. **(b** and **c)** Networks of genes downregulated (b) or upregulated (c) (*P* < 0.01 by Cuffdiff) by feeding relative to fasting, shown as clustered functional categories. *n* = 3 biological replicates of islets pooled from separate mice per group. **(d)** H3K27ac ChIP-seq signal at sites losing or gaining H3K27ac with feeding. *n* = 2 biological replicates of islets pooled from separate mice per group; data from independent ChIP-seq experiments were merged for analysis. Boxplot whiskers span data points within the interquartile range x 1.5. \*\*\**P* < 0.001 by Wilcoxon rank-sum test. **(e** and **f)** Association frequencies between transcription start sites (TSSs) of genes downregulated (e) or upregulated (f) by feeding with the indicated classes of H3K27ac peaks ± 50 kb. Open bars indicate association frequencies expected by chance. \*\*\**P* < 0.001 for enrichment, ^###^*P* < 0.001 for depletion, NS, not significant by permutation test. **(g)** Transcription factor (TF) motifs enriched at sites gaining H3K27ac with feeding relative to all other H3K27ac peaks plotted against mRNA levels of cognate TFs in islets from fed mice. **(h)** Gene set enrichment analysis of genes upregulated in *db/db* compared to control (ctrl, *db/+*) islets (*P* < 0.01 by Cuffdiff; *n* = 3 biological replicates of islets pooled from separate mice per group) against mRNA-seq data from islets after feeding and fasting. **(i)** Log_2_ fold changes in mRNA levels in islets after feeding compared to fasting and in *db/db* compared to control islets. Genes upregulated in both conditions (*P* < 0.01 by Cuffdiff) are highlighted in orange. For clarity, only the upper right quadrant is shown. **(j)** ChIP-seq signal for H3K27ac at the indicated classes of H3K27ac peaks in *db/db* and control islets. *n* = 2 biological replicates of islets pooled from separate mice per group; data from independent ChIP-seq experiments were subsequently merged for analysis. Boxplot whiskers span data points within the interquartile range x 1.5. \*\*\**P* < 0.001 by Wilcoxon rank-sum test; NS, not significant. **(k)** Scatterplot of Lsd1 and H3K27ac ChIP-seq signals at all H3K27ac peaks. Colour intensity of each dot corresponds to Lsd1 ChIP-seq signal intensity. Lsd1 ChIP-seq data are from *n* = 1 biological replicate from pooled islets. Data were highly correlated with Lsd1 ChIP-seq data from an independent biological replicate (see Supplementary Table 3). **(l)** Proportion of Lsd1 peaks associated with the indicated classes of H3K27ac peaks ± 1 kb. Numbers indicate Lsd1 peaks associated with each class of H3K27ac peaks. **(m)** Enrichment test of association frequencies between Lsd1 peaks and the indicated classes of H3K27ac peaks ± 50 kb. Open bars indicate association frequencies expected by chance. \**P* < 0.01 for enrichment, ^#^*P* < 0.01 for depletion by permutation test. **(n)** Lsd1 ChIP-seq signal at the indicated classes of H3K27ac peaks. Boxplot whiskers span data points within the interquartile range x 1.5. \*\*\**P* < 0.001 by Wilcoxon rank-sum test. **(o)** Lsd1 and H3K27ac ChIP-seq genome browser tracks for representative genes exhibiting concordant increases in mRNA and H3K27ac levels with feeding and in *db/db* islets. Source data for all quantifications and exact *P*-values for all indicated statistical tests are provided online.

Metabolic cues have been shown to effect transcriptional changes in part by regulating the epigenome^6,31–33^. In particular, acetylation of histone H3 Lys27 (H3K27ac), a histone modification associated with active promoters and enhancers^34^ (“active chromatin” hereafter), is dynamically regulated in response to changes in nutrient state^6,23,32^. To determine whether islet active chromatin is responsive to nutrient cues, we performed chromatin immunoprecipitation followed by massively parallel DNA sequencing (ChIP-seq) for H3K27ac in islets from fasted and fed mice (**Fig. 1a, Supplementary Table 3**). We found that 48% of H3K27ac peaks exhibited nutrient state-regulated changes in signal (≥ 1.2-fold, *P <* 0.0001, see Methods for details on differential peak calling), the vast majority of which (80%) gained acetylation with feeding (**Fig. 1d**, **Extended Data Fig. 2b, c**, **Supplementary Table 4a**). Feeding-induced deposition of H3K27ac occurred independently of changes in monomethylation of H3 Lys4 (H3K4me1; **Extended Data Fig. 2d**), a histone modification that co-occurs with H3K27ac in active chromatin^34^. Sites gaining H3K27ac with feeding were enriched near genes upregulated in the fed state (**Fig. 1e, f**), suggesting effects of these chromatin changes on gene transcription. Feeding-induced H3K27ac sites were enriched for motifs recognized by ETS and AP-1 family transcription factors (TFs) (**Fig. 1g**, **Supplementary Table 5a**), of which many were expressed in islets (e.g. ETS: *Elk1*, *Elk4*, *Elf1*, *Erg*, *Ets1*, *Etv1*, *Gabpa*; AP-1: *Jun*, *Fos*) (**Fig. 1g**). Notably, these TFs are signal-dependent and have been shown to mediate adaptation to changes in energy state in metabolic tissues^4^. Together, these findings reveal that H3K27ac is dynamically regulated by feeding in islets and suggest signal-dependent regulation of chromatin at these sites.

Adaptive changes in insulin secretion are evoked by both short-term and prolonged changes in nutrient state^16^, the latter being exemplified by compensatory insulin secretion in obesity^17^. During chronic overfeeding, augmented insulin secretion counteracts insulin resistance, thereby preventing hyperglycaemia. To determine whether similar transcriptional and epigenetic changes as observed in our 4-hour-fed versus 16-hour-fast model also occur in response to a sustained increase of nutrient intake, we analysed islets from leptin receptor-deficient *db/db* mice at the onset of hyperglycaemia, when *db/db* islets exhibit insulin hypersecretion indicative of an adaptive response (**Extended Data Fig. 1e, f**). Gene set enrichment analysis (GSEA) revealed significant overlap between genes upregulated in *db/db* islets and genes induced by feeding, including known regulators of nutrient signalling, such as *Irs2*^22,35^, *Mapkapk3*^36^, and *Pak3*^37^ **(Fig. 1h, i, Supplementary Tables 1 and 2b**). This suggests shared adaptive mechanisms in response to short-term or chronic changes in nutrient state. Analysis of chromatin state in *db/db* islets further revealed hyperacetylation of H3K27 and hypermethylation of H3K4 at active chromatin sites, most prominently at feeding-induced H3K27ac sites (**Fig. 1j**, **Extended Data Fig. 2e, f, Supplementary Table 3**). Together, these findings indicate that chronic overfeeding augments nutrient-stimulated histone acetylation and gene transcription concomitant with increased histone methylation.

Dynamics in nutrient state can evoke changes to the epigenome through histone-modifying enzymes that utilize intermediary metabolites as cofactors or substrates or through indirect mechanisms via nutrient-responsive signalling pathways^6,31–33^. Posttranslational histone modifications regulate transcription via recruitment of proteins that impact chromatin accessibility or mediate interactions between regulatory elements. Histone acetylation state exerts a dominant effect on transcription, which is fine-tuned by histone methylation and other modifications^38,39^. One histone-modifying enzyme that has been shown to be associated with acetylated histones and to modulate acetylation and methylation state is the histone demethylase Lsd1^40,41^. In adipose tissue, Lsd1 functions as a signal-dependent chromatin modifier that adapts metabolism to changes in nutrient state^42,43^.

Based on this evidence, we investigated whether Lsd1 associates with nutrient-regulated chromatin in islets. ChIP-seq for Lsd1 in islets revealed a strong positive correlation between Lsd1 and H3K27ac signal intensities (Spearman’s σ = 0.84, *P* < 2.2 x 10^−16^; **Fig. 1k, Supplementary Tables 3 and 6a**), as was observed in other cell types^41,44^. Throughout the genome, Lsd1 predominantly occupied promoters and associated with active and not repressed chromatin (**Extended Data Fig. 2g-k**). Analysis of the extent of overlap between Lsd1-bound sites and feeding-regulated active chromatin revealed enrichment of Lsd1 binding and signal intensity at sites gaining H3K27ac with feeding (**Fig. 1l-n**, **Extended Data Fig. 2k, Supplementary Table 4b**). Compared to feeding-induced H3K27ac regions not bound by Lsd1, H3K27ac regions bound by Lsd1 were enriched for motifs of the ETS and AP-1 TF families, in particular MAPK-dependent Elk TFs (**Extended Data Fig. 2l, Supplementary Table 5b**), suggesting specific association of Lsd1 with sites regulated by signal-dependent TFs. Supporting the relevance of these binding events for gene regulation, Lsd1 peaks were overrepresented near genes upregulated by feeding (**Extended Data Fig. 2m**), such as *Irs2*, *Mapkapk3*, and *Pak3* (**Fig. 1o**, **Extended Data Fig. 2h**). Thus, Lsd1 occupies active chromatin sites that gain acetylation with feeding and associates with feeding-induced genes involved in nutrient signalling.

To investigate the function of Lsd1 in β-cells, we deleted *Lsd1* in β-cells of adult mice using *Pdx1-CreER* (Lsd1^Δβ^ hereafter; **Fig. 2a, b**, **Extended Data Fig. 3a**) and monitored glucose homeostasis. Ad libitum-fed male and female Lsd1^Δβ^ mice began to exhibit hypoglycaemia three weeks after *Lsd1* inactivation (**Fig. 2c**, **Extended Data Fig. 3b**). The same phenotype was observed following *Lsd1* deletion with the *MIP-CreER* transgene (**Extended Data Fig. 3c**). Lsd1^Δβ^ and control mice did not differ with regard to food consumption or body weight, and insulin sensitivity was modestly reduced (**Extended Data Fig. 3d-f**), excluding reduced caloric intake or increased insulin sensitivity as the primary cause of hypoglycaemia. Analysis of β-cell mass, islet endocrine cell type composition, and pancreatic insulin content (**Extended Data Fig. 3g-i**) further ruled out β-cell hyperplasia as underlying the hypoglycaemia.

**Figure 2.**
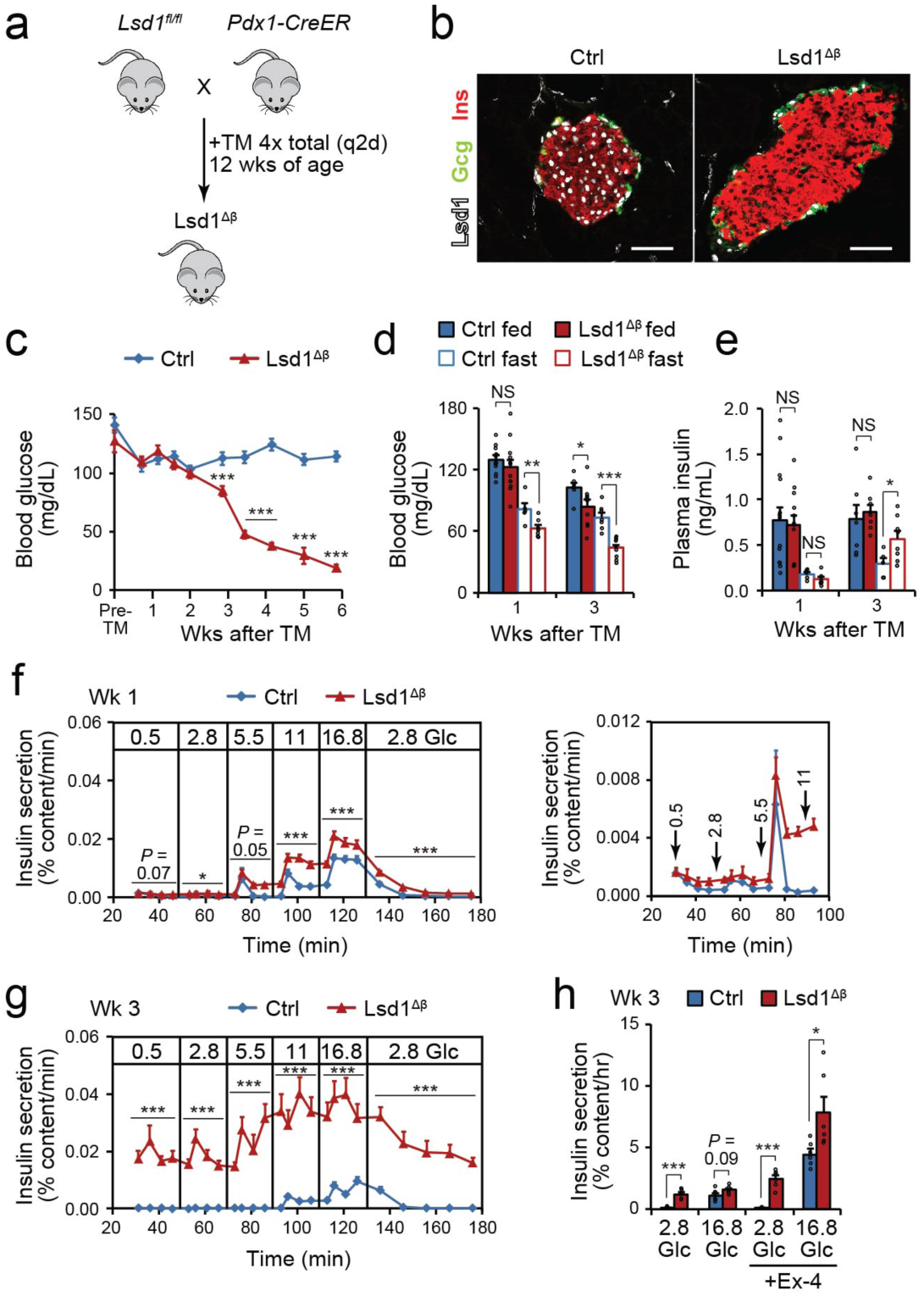
*Lsd1* inactivation in β-cells causes insulin hypersecretion and hypoglycaemia. **(a)** Schematic of alleles and treatments used to inactivate *Lsd1*. TM, tamoxifen; wks, weeks; q2d, every other day. **(b)** Immunofluorescence staining for insulin (Ins), glucagon (Gcg) and Lsd1 on pancreas sections from control (ctrl, control TM-treated *Lsd1^fl/+^*; *Pdx1-CreER* mice) and Lsd1^Δβ^ mice two days after TM treatment. Scale bar, 50 µm. **(c)** Time course of ad libitum-fed blood glucose levels in control (TM-treated *Lsd1^+/+^*; *Pdx1-CreER* mice; *n* = 9) and Lsd1^Δβ^ (*n* = 10) mice. Pre-TM, within 3 days prior to initial TM injection. \*\*\**P* < 0.001 by unpaired two-tailed t-test. **(d** and **e)** Blood glucose (d) and serum insulin (e) levels in ad libitum-fed and 16 hour-fasted mice of the indicated genotypes. Control fed week one: *n =* 15, Lsd1^Δβ^ fed week one: *n =* 13, control fasted week one: *n =* 5, Lsd1^Δβ^ fasted week one: *n =* 8, control fed week three and control fasted week three: *n =* 7, all other groups, *n =* 9 mice (d). Control fed week one: *n =* 15, Lsd1^Δβ^ fed week one: *n =* 12, control fasted week one and control fasted week three: *n =* 5, Lsd1^Δβ^ fed week three and Lsd1^Δβ^ fasted week three: *n =* 8, all other groups: *n =* 7 mice (e). \**P* < 0.05, \*\**P* < 0.01, \*\*\**P* < 0.001 by unpaired two-tailed t-test with Welch’s correction for unequal variance as necessary. **(f** and **g)** Insulin secretion by control and Lsd1^Δβ^ islets during perifusion with the indicated glucose (Glc) concentrations (in mM) at one week (f) and three weeks (g) following TM treatment (*n* = 4 pools of 130 islets each). Right (f), data shown at a reduced scale for the indicated time points. \**P* < 0.05, \*\*\**P* < 0.001 by two-way ANOVA for genotype for each time block. **(h)** Static insulin secretion assay in control and Lsd1^Δβ^ islets stimulated with the indicated glucose (Glc) concentrations (in mM) with and without Exendin-4 (Ex-4) three weeks following TM treatment. Control 2.8 mM glucose: *n* = 4 pools of 10 islets each, control 2.8 mM glucose + Ex-4: *n* = 5 islet pools, all other groups: *n* = 6 islet pools. \**P* < 0.05, \*\*\**P* < 0.001 by unpaired two-tailed t-test with Welch’s correction for unequal variance as necessary. Data are shown as mean ± S.E.M. Source data for all quantifications and exact *P*-values for all indicated statistical tests are provided online.

To investigate the progressive nature and possible nutrient dependency of the hypoglycaemic phenotype, we analysed blood glucose levels in Lsd1^Δβ^ mice in different nutritional states before and at the onset of overt hypoglycaemia. One week after *Lsd1* deletion, ad libitum-fed mice maintained normal blood glucose levels throughout the day (**Extended Data Fig. 3j**). An intraperitoneal glucose tolerance test further indicated that glucose sensing during stimulation is normal one week following *Lsd1* inactivation (**Extended Data Fig. 3k**). However, after overnight fasting Lsd1^Δβ^ mice exhibited significantly lower blood glucose levels at this time point (**Fig. 2d**). Fasting hypoglycaemia was accompanied by inappropriately high plasma insulin levels in the fasted but not the fed state (**Fig. 2e**). It is unlikely that this phenotype is caused by a defective counter-regulatory response because blood glucagon levels were not reduced in fasted Lsd1^Δβ^ mice (**Extended Data Fig. 3l**). These findings suggest that adaptation of the β-cell insulin secretory response to fasting is disrupted in Lsd1^Δβ^ mice.

To test whether Lsd1^Δβ^ mice exhibit insulin hypersecretion at sub-stimulatory glucose concentrations, we studied Lsd1^Δβ^ islets in perifusion experiments. One week after *Lsd1* deletion, Lsd1^Δβ^ islets exhibited increased insulin secretion at intermediate and high glucose concentrations compared to control islets (**Fig. 2f**), and by the three-week time point, islets secreted insulin even at supraphysiologically low glucose levels (**Fig. 2g**). Thus, Lsd1^Δβ^ islets hypersecrete insulin in response to glucose prior to an overt in vivo phenotype, suggesting transient in vivo compensation. These results demonstrate that *Lsd1* inactivation results in a cell-autonomous increase of GSIS and failure to suppress insulin secretion in response to hypoglycaemia.

Insulin secretion is modulated by fatty acids and amino acids as well as hormones^9,11^; therefore, we tested whether islets from Lsd1^Δβ^ mice respond aberrantly to other secretagogues. While Lsd1^Δβ^ islets responded normally to the fatty acid palmitate and to co-stimulation with leucine and glutamine (**Extended Data Fig. 4a**), they exhibited an exaggerated response to Exendin-4, an analogue of the feeding-induced hormone Glp-1 (**Fig. 2h**). The increased sensitivity to Glp-1 or other cAMP-generating pathways^7,8^ could contribute to hypoglycaemia in the fed state following *Lsd1* inactivation (**Fig. 2d**).

Glucose sensing by the β-cell relies on glucose metabolism, such that the rate of glycolysis is determined by blood glucose concentration. An increase in the β-cell ATP/ADP ratio in response to glucose triggers insulin secretion by inducing closure of an ATP-sensitive potassium (K_ATP_) channel, membrane depolarization, and subsequent calcium (Ca^2+^) influx^12^. Having found that insulin secretion becomes progressively uncoupled from glucose levels in *Lsd1*-deficient β-cells, we sought to determine which step(s) of stimulus-secretion coupling is deregulated. As a readout for metabolic activity, we measured respiration under basal and stimulatory glucose conditions (2.8 mM and 16.8 mM glucose, respectively), and found that Lsd1^Δβ^ islets consumed more oxygen in low glucose than control islets (**Fig. 3a**). Reduced respiration following inhibition of ATP synthase by oligomycin indicates that oxygen consumption in Lsd1^Δβ^ islets is coupled to ATP synthesis (**Fig. 3a**). Accordingly, ATP content in low glucose was elevated in Lsd1^Δβ^ islets (**Fig. 3b**). To study stimulus-secretion coupling downstream of ATP production in Lsd1^Δβ^ islets, we measured Ca^2+^ influx in basal and stimulatory glucose concentrations. Consistent with their higher ATP content, *Lsd1*-deficient β-cells exhibited elevated Ca^2+^ levels in basal glucose levels akin to control β-cells under stimulatory glucose concentrations (**Fig. 3c**), suggesting constitutive activation of voltage-gated Ca^2+^ channels. To test whether Ca^2+^ influx in Lsd1^Δβ^ β-cells is triggered by tonic inhibition of K_ATP_ channels in basal glucose concentrations, we treated islets with the K_ATP_ channel opener diazoxide. Showing the K_ATP_ channel-dependency of Ca^2+^ influx in *Lsd1*-deficient β-cells, diazoxide inhibited Ca^2+^ influx (**Fig. 3d**) and reduced basal insulin secretion (**Fig. 3e**). Together, these results suggest that altered metabolism in Lsd1^Δβ^ islets promotes basal insulin hypersecretion via closure of K_ATP_ channels and subsequent activation of voltage-gated Ca^2+^ channels.

**Figure 3.**
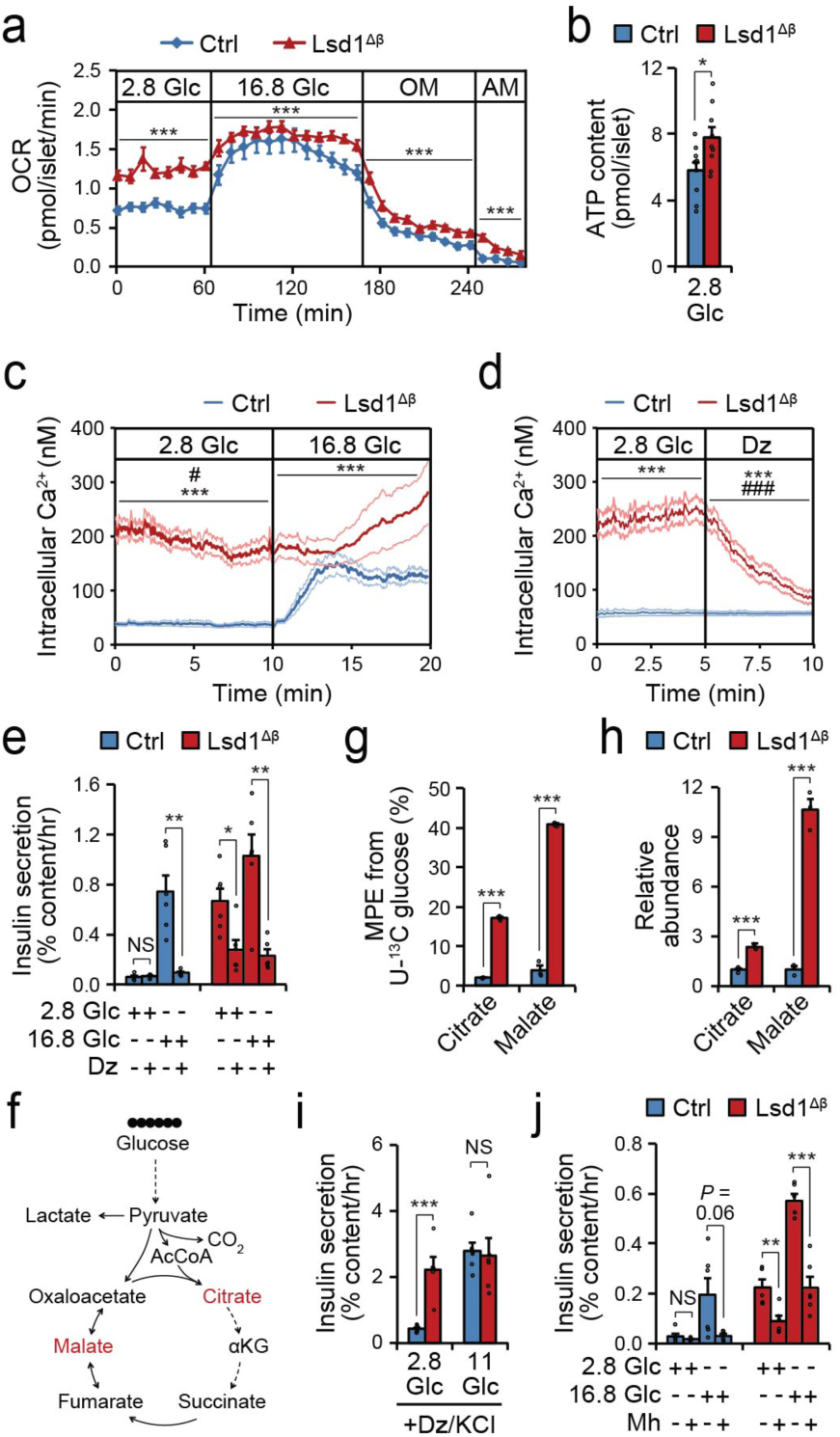
Accelerated glucose metabolism promotes insulin hypersecretion in Lsd1^Δβ^ islets. **(a)** Oxygen consumption rate (OCR) of control (ctrl) and Lsd1^Δβ^ islets treated sequentially with the indicated glucose (Glc) concentrations (in mM), oligomycin (OM), and antimycin A (AM). Control: *n* = 10 pools of 60 islets each, Lsd1^Δβ^: *n* = 8 islet pools. \*\*\**P* < 0.001 by two-way ANOVA for genotype for each time block. **(b)** ATP content of control and Lsd1^Δβ^ islets cultured in 2.8 mM glucose. Control: *n* = 10 pools of 20 islets each, Lsd1^Δβ^: *n* = 9 islet pools. \**P* < 0.05 by unpaired two-tailed t-test. **(c** and **d)** Intracellular Ca^2+^ concentration in β-cells from control and Lsd1^Δβ^ mice cultured in the indicated glucose (Glc) concentrations (in mM) (c) or treated with the K_ATP_ channel opener diazoxide (Dz) (d). Control: *n* = 38 (c) or 46 (d) β-cells, Lsd1^Δβ^: *n* = 32 (c) or 36 (d) β-cells. Results are representative of 3 independent experiments. \*\*\**P* < 0.001 by two-way ANOVA for genotype for each time block. #*P* < 0.05, ###*P* < 0.001 by two-way ANOVA for the interaction between genotype and time for each time block. **(e)** Static insulin secretion assay in control and Lsd1^Δβ^ islets stimulated with the indicated glucose concentrations with and without Dz. Control 2.8 mM glucose: *n* = 5 pools of 10 islets each, all other groups: *n* = 6 islet pools. \**P* < 0.05, \*\**P* < 0.01 by unpaired two-tailed t-test with Welch’s correction for unequal variance as necessary. NS, not significant. **(f)** Schematic of isotopic glucose tracing experiment. Metabolites displayed in (g, h) are indicated in red. **(g** and **h)** Molar percent enrichment (MPE) of ^13^C (g) and relative abundances (h) of the indicated metabolites following two hours of tracing in 2.8 mM U-^13^C glucose. *n* = 3 pools of 220 islets each. \*\*\**P* < 0.001 by unpaired two-tailed t-test. **(i** and **j)** Static insulin secretion assay of control and Lsd1^Δβ^ islets stimulated with the indicated glucose concentrations under depolarizing conditions (30 mM KCl and 100 µM Dz) (i) or with and without the glycolysis inhibitor mannoheptulose (Mh) (j). Lsd1^Δβ^ 2.8 mM glucose/Dz/KCl (i), Lsd1^Δβ^ 2.8 mM glucose (j), and control 2.8 mM glucose/Mh (j): *n* = 5 pools of 10 islets each, all other groups: *n* = 6 islet pools. \*\**P* < 0.01, \*\*\**P* < 0.001 by unpaired two-tailed t-test with Welch’s correction for unequal variance as necessary. NS, not significant. Islets were isolated from control and Lsd1^Δβ^ animals three weeks following tamoxifen treatment for all experiments. Data are shown as mean ± S.E.M. Source data for all quantifications and exact *P*-values for all indicated statistical tests are provided online.

To directly test whether the metabolism of glucose fuels ATP production and constitutive insulin secretion in Lsd1^Δβ^ islets, we determined the metabolic fate of glucose in islets using isotopic tracing of uniformly-^13^C-labeled glucose (hereafter U-^13^C-glucose; **Fig. 3f**). For these experiments, control and Lsd1^Δβ^ islets were incubated in 2.8 mM of U-^13^C-glucose, reflective of a non-stimulatory glucose concentration, and analysed by targeted metabolomics to determine abundance and isotopic labelling of glucose-derived metabolites. Lsd1^Δβ^ islets exhibited increased isotopic labelling and accumulation of citrate and malate (**Fig. 3g, h**), which have been shown to increase during glucose stimulation^45,46^. Lsd1^Δβ^ islets also exhibited increased ^13^C labelling of glycine, aspartate, and glutamate (**Extended Data Fig. 4b**), reflecting accelerated glucose metabolism via glycolysis and the TCA cycle. Lactate production was not altered (**Extended Data Fig 4c**), suggesting that Lsd1^Δβ^ islets metabolize much of this glucose in mitochondria. These results indicate that *Lsd1* deficiency gives rise to a basal metabolic state that resembles the glucose-stimulated state of normal islets.

Glucose metabolism stimulates insulin secretion via both “triggering” and “amplifying” pathways^12^. The triggering pathway is stimulated when ATP produced from glucose oxidation inhibits K_ATP_ channels, resulting in depolarization and opening of voltage-gated Ca^2+^ channels. The amplifying pathway potentiates the effect of Ca^2+^ upon insulin vesicle exocytosis, which occurs in part via accumulation of intermediary metabolites^9,46^. Having observed augmented production of glucose-derived citrate and malate in Lsd1^Δβ^ islets (**Fig. 3g, h**), we predicted that these metabolites could hyperactivate the amplifying pathway. Thus, we assessed activity of the amplifying pathway by constitutively activating the triggering pathway with KCl and preventing metabolic effects on the K_ATP_ channel with simultaneous diazoxide treatment. This revealed constitutive activation of the amplifying pathway in Lsd1^Δβ^ islets (**Fig 3i**). Supporting the important role of accelerated glucose metabolism in basal insulin hypersecretion by *Lsd1*-deficient β-cells, inhibition of glycolysis with mannoheptulose partially rescued basal insulin hypersecretion by Lsd1^Δβ^ islets (**Fig. 3j**). Altogether, these findings suggest that aberrant glucose metabolism in the unstimulated state activates both triggering and amplifying pathways of insulin secretion in Lsd1^Δβ^ islets. Dual activation of these pathways likely underlies the increased severity of the phenotype compared to models of triggering pathway deregulation such as *Sur1* inactivation^47^.

Having observed uncoupling of insulin secretion from nutrient state in Lsd1^Δβ^ mice, we sought to determine the molecular mechanisms underlying these changes. To this end, we characterized transcriptomes and epigenomes of Lsd1^Δβ^ islets before and at the onset of hypoglycaemia at one and three weeks after *Lsd1* deletion, respectively (**Fig. 4a**). mRNA-seq analysis revealed that many transcriptional changes observed three weeks after *Lsd1* inactivation were already present at week one (280 of 643 genes upregulated at week three and 124 of 350 genes downregulated at week three; *P* < 0.01 using Cuffdiff; Spearman’s σ = 0.49, *P* < 2.2 x 10^−16^ for regulated genes; **Fig. 4b**, **Supplementary Tables 1 and 2c**). Thus, loss of *Lsd1* in β-cells has effects on gene transcription prior to manifestation of overt hypoglycaemia. Concordant with the transcriptomic changes, Lsd1^Δβ^ islets also exhibited early alterations in islet chromatin state. Of 16,324 total differential H3K27ac peaks (≥ 1.5 absolute fold difference) between Lsd1^Δβ^ and control islets at either one or three weeks following *Lsd1* inactivation, 3,076 peaks (19% of total differential peaks) were already changed at week one (**Fig. 4c**, **Extended Data Fig. 5a**, **Supplementary Tables 3 and 4c**). Remarkably, early H3K27ac changes comprised mostly a gain in acetylation (3,329 differential peaks gaining vs. 49 losing acetylation at week one). Amongst all classes of H3K27ac sites, early hyperacetylated sites exhibited the strongest Lsd1 binding signal (**Fig. 4d**), suggesting direct effects of Lsd1. These sites also exhibited H3K4me1 accumulation (**Extended Data Fig. 5b-d**, **Supplementary Table 4d**), consistent with Lsd1’s activity as a H3K4 mono- and di-demethylase^48^ and further supporting a direct role for Lsd1 at these sites.

**Figure 4.**
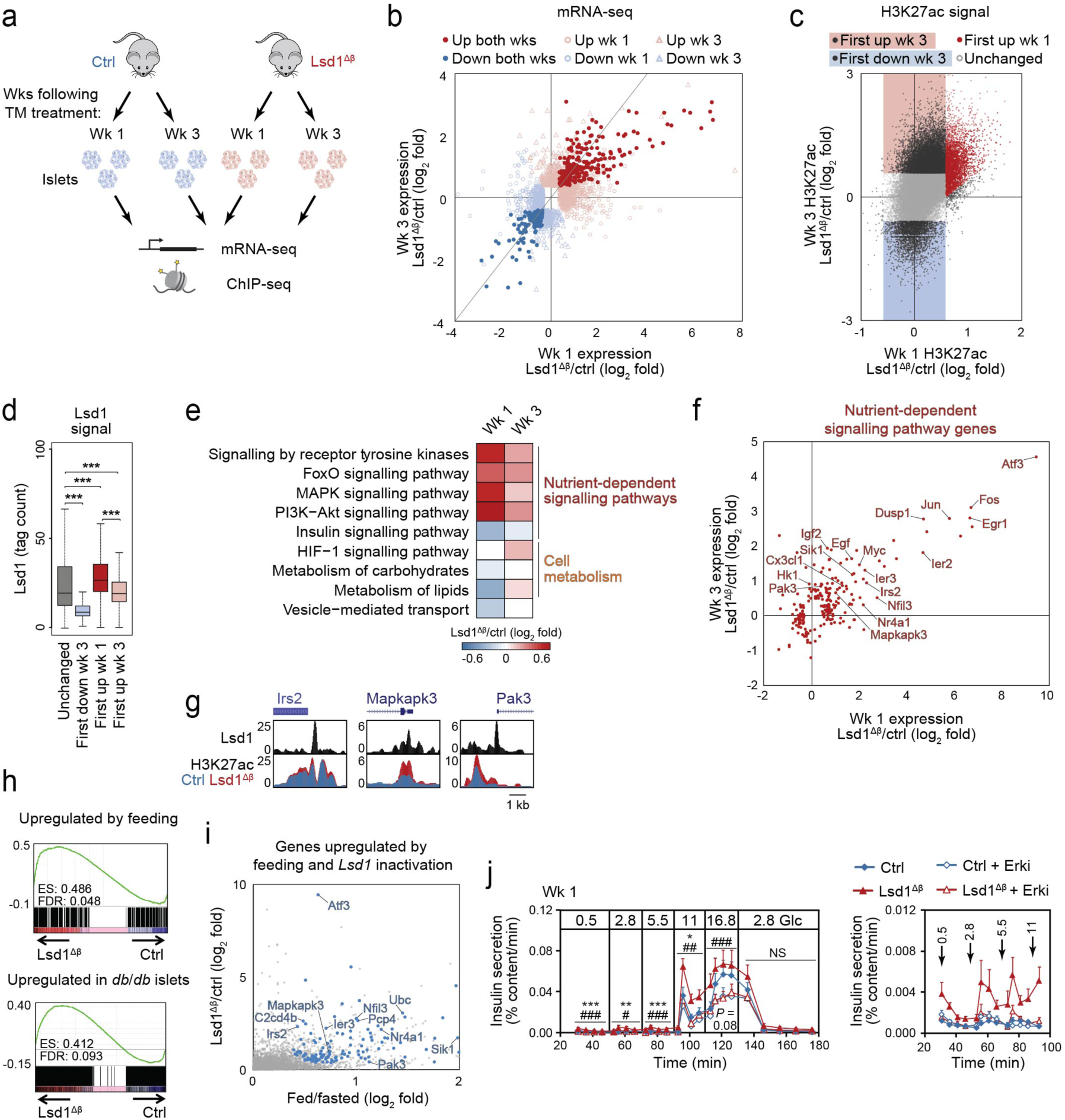
*Lsd1* inactivation in β-cells deregulates genes involved in nutrient-dependent signalling and cell metabolism. **(a)** Schematic of experiments performed. Ctrl, control; TM, tamoxifen; wk/s, week/s. **(b)** Relative mRNA levels for differentially expressed genes (*P* < 0.01 by Cuffdiff) in Lsd1^Δβ^ islets compared to control islets at the indicated time points following TM treatment. The grey line indicating a slope of 1 provides a reference for corresponding mRNA levels between time points. Lsd1^Δβ^ week three: *n* = 5 biological replicates of islets pooled from separate mice per group, all other groups: *n* = 3 biological replicates. **(c)** H3K27ac ChIP-seq signal at H3K27ac peaks (from Extended Data Fig. 2b) plotted as relative tag density in Lsd1^Δβ^ islets compared to control islets at the indicated time points following TM treatment. Peaks were classified by the timepoint when relative tag density in Lsd1^Δβ^ islets exceeded 1.5-absolute fold difference compared to control islets (see Methods for additional details on peak classification). *n* = 2 biological replicates of islets pooled from separate mice per group; data from independent ChIP-seq experiments were subsequently merged for analysis. **(d)** Lsd1 ChIP-seq signal at the indicated classes of H3K27ac peaks (from c). Lsd1 ChIP-seq data are from *n* = 1 biological replicate from pooled islets. Data were validated to be highly correlated with ChIP-seq data prepared from an independent biological replicate. Boxplot whiskers span data points within the interquartile range x 1.5. \*\*\**P* < 0.001 by Wilcoxon rank-sum test. **(e)** Median log_2_ fold expression changes in mRNA levels between Lsd1^Δβ^ and control islets within each of the indicated functional categories at one and three weeks following TM treatment. **(f)** Log_2_ fold changes in mRNA levels between Lsd1^Δβ^ and control islets at one and three weeks following TM treatment. Genes annotated to nutrient-dependent signalling pathways (from e) and differentially expressed (*P* < 0.01 by Cuffdiff) in Lsd1^Δβ^ islets compared to control islets at either one or three weeks following TM treatment are plotted. **(g)** Lsd1 and H3K27ac ChIP-seq genome browser tracks at representative nutrient-dependent signalling pathway genes in control and Lsd1^Δβ^ islets one week following TM treatment. **(h)** Gene set enrichment analysis of genes upregulated by feeding in islets (top, *P* < 0.01 by Cuffdiff) and in *db/db* compared to control (*db/+*) islets (bottom, *P* < 0.01 by Cuffdiff) against mRNA-seq data from Lsd1^Δβ^ and control islets one week following TM treatment. **(i)** Log_2_ fold changes in mRNA levels in islets after feeding compared to fasting and in Lsd1^Δβ^ compared to control islets one week following TM treatment. Genes upregulated in both conditions (*P* < 0.01 by Cuffdiff) are highlighted in blue. For clarity, only the upper right quadrant is shown. **(j)** Insulin secretion by control and Lsd1^Δβ^ islets during perifusion with the indicated glucose (Glc) concentrations (in mM) following 24-hour treatment with SCH772984 (Erki) or vehicle. Islets were isolated one week following TM treatment. Right, data shown at a reduced scale for the indicated time points. *n* = 6 pools of 130 islets each. Data in are shown as mean ± S.E.M. \**P* < 0.05, \*\**P* < 0.01, \*\*\**P* < 0.001 vehicle-treated Lsd1^Δβ^ relative to vehicle-treated control islets by two-way ANOVA for genotype for each time block, #*P* < 0.05, ##*P* < 0.01, ###*P* < 0.001 Erki-treated relative to vehicle-treated Lsd1^Δβ^ islets by two-way ANOVA for treatment group for each time block. Source data for all quantifications and exact *P*-values for all indicated statistical tests are provided online.

The similarities between effects of *Lsd1* deletion (**Fig. 4c**), short-term feeding (**Fig. 1d**, **Extended Data Fig. 2b, c**), and chronic overfeeding (**Fig. 1j**) on islet H3K27ac raised the possibility that Lsd1 dampens acetylation at nutrient-regulated H3K27ac sites. Indeed, sites that gained H3K27ac with feeding exhibited excess H3K27ac deposition in Lsd1^Δβ^ islets (**Extended Data Fig. 5e**). Uncoupling of histone acetylation from feeding state in Lsd1^Δβ^ islets indicates that Lsd1 is required for the β-cell to interpret nutrient signals at the level of the epigenome. However, as in *db/db* islets (**Extended Data Fig. 2e**), hyperacetylation in Lsd1^Δβ^ islets was also evident at H3K27ac sites not regulated by short-term feeding (**Extended Data Fig. 5e**). Similarly, Lsd1^Δβ^ (**Extended Data Fig. 5f, g**) and *db/db* islets (**Extended Data Fig. 2f, h**) exhibited increased H3K4 monomethylation at feeding-induced H3K27ac sites, whereas a gain in H3K4me1 was not observed after short-term feeding (**Extended Data Fig. 2d, h**). Thus, *Lsd1* deletion or prolonged overfeeding have more pronounced effects on the epigenome relative to short-term feeding.

To identify gene expression changes that drive insulin hypersecretion following *Lsd1* inactivation, we determined functional category enrichment among differentially expressed genes in Lsd1^Δβ^ islets. Enrichments were independently assessed for genes regulated at one or three weeks after *Lsd1* inactivation to account for transient or delayed regulation (**Supplementary Tables 2c and 7a-c**). This analysis revealed deregulation of processes involved in the control of insulin secretion, such as nutrient-dependent signalling, cell metabolism, and vesicle transport, with most genes exhibiting concordant changes at both time points (**Fig. 4e, f**, **Extended Data Fig. 6a, b**). Among the deregulated genes were ligands that stimulate receptors of nutrient-responsive signalling pathways in β-cells, such as *Cx3cl1*, *Egf*, and *Igf2*, which have positive effects on insulin secretion^49–51^. Additional deregulated genes included *Irs2*, *Dusp1*, *Mapkapk3*, and *Pak3,* which are intermediates of the PI3K/Akt and MAPK signalling pathways that are known to promote insulin secretion^22,35,52,53^. Of interest was the high expression of *Atf3*, *Fos, Jun,* and *Nr4a1* in Lsd1^Δβ^ islets. These TFs are stimulus-induced immediate-early genes, which are transcribed within minutes after activation of a signalling pathway^21,54^. Nr4a1, Atf3, and Fos have been shown to regulate insulin secretion in β-cells^55–57^, suggesting that immediate-early genes could mediate adaptation of the insulin secretory response to changes in the nutrient environment. Transcription start sites (TSSs) of many nutrient response genes, such as *Irs2*, *Mapkapk3*, and *Pak3* (**Fig. 4g**, **Extended Data Fig. 5g**), were Lsd1-bound and exhibited H3K27 hyperacetylation and increased H3K4 monomethylation in Lsd1^Δβ^ islets, indicating direct effects of Lsd1 on chromatin state at these gene loci. Of note, the low-K_m_ glucose-phosphorylating enzyme hexokinase 1 (*Hk1*) was only upregulated at three weeks but not at one week after *Lsd1* inactivation (**Fig. 4f**). Hk1 is normally absent from β-cells^58^, which enables β-cell glucose sensing through the high-K_m_ glucose-phosphorylating enzyme glucokinase (Gck). Ectopic expression of Hk1 in β-cells accelerates glycolysis in low glucose^59^ and causes basal insulin hypersecretion and hypoglycaemia^60^ similar to Lsd1^Δβ^ mice three weeks after *Lsd1* inactivation (**Fig. 2d, g**). This suggests that the delayed upregulation of Hk1 could be a major contributor to the late onset of overt hypoglycaemia in Lsd1^Δβ^ mice. An additional driver for progressive dysregulation of insulin secretion could be genes involved in HIF-1α signalling, which were similarly first increased three weeks following *Lsd1* deletion (**Fig. 4e**). HIF-1α mediates β-cell metabolic reprogramming and is induced by chronic accumulation of cAMP, a second messenger of Glp-1 and glucagon receptor signalling^61,62^. Taken together, these observations support a model whereby *Lsd1* inactivation deregulates nutrient-dependent signalling pathways and leads to metabolic reprogramming of β-cells, thereby uncoupling insulin secretion from nutrient state.

The overlap in cellular processes affected by *Lsd1* deletion (**Fig. 4e**) and induced by feeding (**Fig. 1c**) led us to examine whether *Lsd1* inactivation deregulates similar genes as those regulated during adaptation to changing nutrient states (e.g. *Irs2*, *Mapkapk3*, *Pak3*, *Atf3*, *Nr4a1*, and *Nr4a2*). GSEA revealed enrichment of genes upregulated by feeding as well as genes upregulated in *db/db* islets among those overexpressed in Lsd1^Δβ^ islets (**Fig. 4h, i**, **Extended Data Fig. 7a**). Similar genes were acutely induced in β-cells^21^ and islets by co-stimulation with glucose and the Glp-1 effector cAMP (**Extended Data Fig. 7b-d**), indicating that *Lsd1*-deficient β-cells resemble β-cells in a nutrient-stimulated state despite being exposed to similar nutrient environments in vivo at one week after *Lsd1* inactivation (**Extended Data Fig. 3j, k**). Together, these findings show that Lsd1 mediates coupling of environmental nutrient signals to transcription of genes associated with β-cell functional adaptation.

The prevalence of nutrient response genes with known roles in insulin secretion (e.g. *Atf3*, *Irs2*, and *Nr4a1*)^35,55,56^ among Lsd1-regulated transcripts (**Fig. 4i**) suggested that Lsd1 could adapt the insulin secretory response by modulating nutrient-dependent signalling. Several lines of evidence pointed to an important role for MAPK signalling in this process. First, nutrient stimulation increases ERK phosphorylation in β-cells^26^. Furthermore, both feeding and *Lsd1* deletion stimulated expression of MAPK pathway-associated genes (**Fig. 1c**, **Fig. 4e**). Finally, feeding-regulated H3K27ac sites bound by Lsd1 were highly enriched for recognition motifs of the MAPK effectors Elk1 and Elk4 (**Extended Data Fig. 2l**). To determine the importance of MAPK signalling for insulin hypersecretion in Lsd1^Δβ^ islets, we treated control and Lsd1^Δβ^ islets with an ERK inhibitor for 24 hours. Indeed, ERK inhibitor treatment rendered the insulin secretory response of Lsd1^Δβ^ islets indistinguishable from control islets (**Fig. 4j**). Together, these results demonstrate that inhibition of MAPK signalling reverses insulin hypersecretion induced by *Lsd1* inactivation, indicating functional convergence of Lsd1-regulated programs and the MAPK signalling pathway in β-cells.

To next determine whether LSD1 function is conserved in human β-cells, we treated cadaveric human islets with the LSD1 inhibitor tranylcypromine (LSD1i) or conducted knockdown in human β-cells and performed GSIS assays. Similar to Lsd1^Δβ^ murine islets (**Fig. 2h**), LSD1i-treated human islets or β-cells transduced with an *LSD1*-shRNA exhibited increased basal insulin secretion (**Fig. 5a**, **Extended Data Fig. 8a-d**). Transcriptome analysis of LSD1i-treated human islets revealed that genes induced by LSD1i treatment were also upregulated in Lsd1^Δβ^ mouse islets (**Fig. 5b**), including immediate-early genes (*ATF3*, *EGR1, IER3, JUN, JUND, MAFF, KLF4, NFIL3*) and the nutrient-response gene *IRS2* (**Supplementary Tables 1 and 2d**).

**Figure 5.**
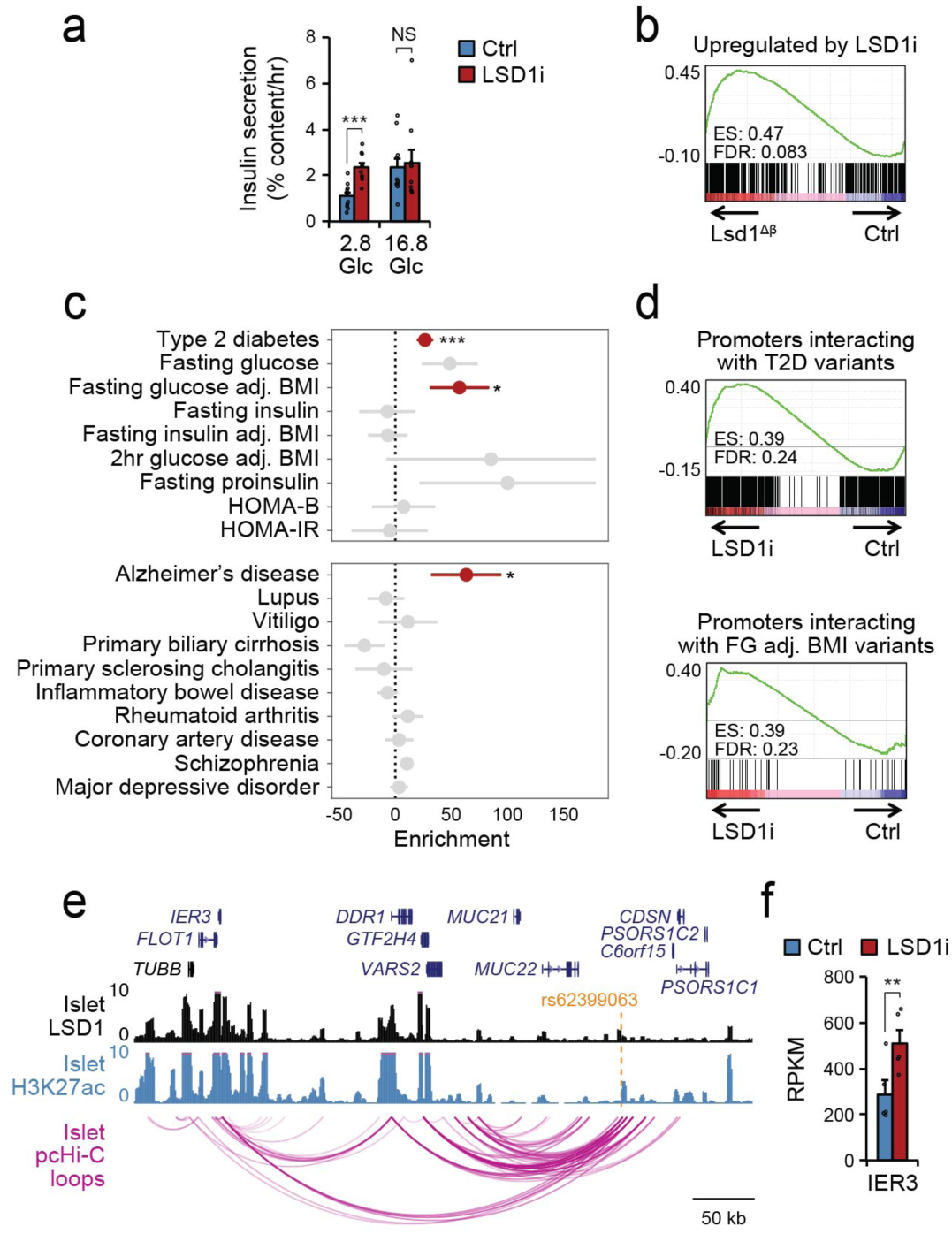
LSD1 regulates insulin secretion in human islets and its binding sites are enriched for type 2 diabetes-associated variants. **(a)** Static insulin secretion assay in human islets stimulated with the indicated glucose (Glc) concentrations (in mM) following 24-hour treatment with the LSD1 inhibitor (LSD1i) TCP or vehicle control (ctrl). *n* = 8 donors. Data are shown as mean ± S.E.M. \*\*\**P* < 0.001 by paired two-tailed t-test. **(b)** Gene set enrichment analysis of genes upregulated by LSD1i treatment in human islets (*P* < 0.01 by Cuffdiff; *n* = 5 donors each treated with vehicle or LSD1i) against mRNA-seq data from Lsd1^Δβ^ and control mouse islets one week following TM treatment. LSD1i-regulated genes were converted to mouse orthologues for enrichment analysis. **(c)** GWAS enrichment (%h^2^/%SNPs) of metabolic (top) and non-metabolic traits (bottom) at LSD1 ChIP-seq peaks in human islets using LD-score regression. Data shown represent enrichment estimate and standard error, and significant estimates are highlighted in red. *** P < 0.001, * P < 0.05 by LD-score regression. Data from independent ChIP-seq experiments from 2 donors were merged for analysis. **(d)** Gene set enrichment analysis of genes whose promoters interact with an LSD1-bound site containing a type 2 diabetes (T2D)-associated variant (top) or a BMI-adjusted fasting glucose (FG adj. BMI)-associated variant (bottom) from promoter capture Hi-C data against mRNA-seq data from LSD1i-treated and control human islets. **(e)** LSD1 and H3K27ac ChIP-seq and promoter capture Hi-C (pcHi-C)^23^ genome browser tracks showing interaction between the LSD1-bound site containing T2D-associated variant rs62399063 (highlighted in orange) and the *IER3* promoter in human islets. **(f)** Bar graph of *IER3* mRNA level in human islets treated with LSD1i as in (a). Data shown as mean ± S.E.M. \*\**P* < 0.01 by Cuffdiff. Source data for all quantifications and exact *P*-values for all indicated statistical tests are provided online.

To further determine whether LSD1 associates with active chromatin in human islets as observed in mice, we conducted ChIP-seq for LSD1 (**Supplementary Tables 3 and 6b**). As in mouse islets (**Fig. 1k, l**, **Extended Data Fig. 2g, i**), LSD1 predominantly occupied active promoters and enhancers in human islets (**Extended Data Fig. 8e, Supplementary Table 6b**), with LSD1 and H3K27ac signal intensities correlating across the genome (Spearman’s σ = 0.79, *P* < 2.2 x 10^−16^; **Extended Data Fig. 8f-h, Supplementary Table 6c**).

Given the here-demonstrated role for LSD1 in the regulation of insulin secretion, we postulated that genetic variants associated with traits relevant to insulin secretion could exert their function through sites occupied by LSD1. To test this prediction, we calculated enrichment of genetic variants at LSD1-bound sites for association with type 2 diabetes (T2D)^63^, diabetes-related quantitative phenotypes^64–67^, and other complex traits and diseases for calibration^68–77^. We observed significant enrichment (*P* < 0.05) of body mass index-adjusted fasting glucose (FG adj. BMI) and T2D association (FG adj. BMI, enrichment = 57.2, *P* = 0.03; T2D, enrichment = 26.5, *P* = 4.65 x 10^−4^) (**Fig. 5c**). As expected, traits associated with insulin resistance (e.g. fasting insulin) as well as most non-metabolic traits showed no evidence for enrichment. The one exception was Alzheimer’s disease, which could indicate that LSD1 regulates similar genes in brain as in islets, consistent with known functions for Lsd1 in neuronal gene regulation^78^.

In order to define the LSD1-mediated gene networks affected by T2D- and fasting glucose-associated variation, we annotated genetic variants in LSD1-bound sites with evidence for T2D or FG adj. BMI association with their putative target genes in islets using chromatin loops derived from promoter capture Hi-C (pcHi-C) data^23^ (**Supplementary Table 8a-b**). We observed significant enrichment of genes looped to FG adj. BMI and T2D-associated LSD1-bound sites among genes upregulated in LSD1i-treated islets (**Fig. 5d**). For example, at loci harbouring the signal-dependent immediate-early genes *IER3* and *JUN*, T2D-associated variants in LSD1-occupied enhancers exhibited chromatin looping to the *IER3* and *JUN* promoters, respectively (**Fig. 5e**, **Extended Data Fig. 8i**). IER3 and JUN mRNAs were both upregulated by *LSD1* inactivation in human and mouse islets (**Fig. 4f**, **Fig. 5f**, **Extended Data Fig. 8j**). In sum, our findings identify a key role for LSD1 in preserving glucose homeostasis of mice and humans by ensuring the insulin secretory response is coupled to nutrient state.

## Discussion

Through integrated studies of β-cell physiology, epigenomics, and transcriptomics, we have established nutrient-dependent regulation of the islet epigenome as a mechanism for β-cell functional adaptation to changes in organismal insulin demand. We find that feeding induces histone hyperacetylation and transcription at gene loci involved in the regulation of β-cell nutrient signalling and metabolism, indicating that nutrients sensitize the response of β-cells to insulin secretory cues through effects on the epigenome. Our analysis identifies Lsd1 as a component of the transcriptional complexes residing at sites whose acetylation levels are coupled to nutrient state. By dampening histone acetylation and methylation at these sites, Lsd1 prevents aberrantly high expression of nutrient-induced genes, thereby counteracting the sensitisation of the β-cell insulin secretory response by nutrient cues. The function of Lsd1 in β-cells has similarity with its role in neurons, where Lsd1 regulates chromatin state and expression of genes that are modulated by excitatory stimuli^78^. Lsd1 has been shown to affect histone acetylation via cross-talk with histone methylation^40^, demethylation of TFs^79^, and through non-enzymatic scaffolding functions^80^. Further studies will be needed to define the precise mechanisms through which Lsd1 regulates nutrient-responsive chromatin in β-cells.

Our findings suggest that regulation of chromatin state at active regulatory elements of genes involved in metabolism and nutrient signalling provides a mechanism for β-cells to adapt their insulin secretory response to changing insulin demands. At nutrient-responsive genes, we observed qualitatively similar changes in H3K27 acetylation and gene expression in a short-term feeding model as in the *db/db* model of chronic overnutrition at a time point when *db/db* islets exhibit adaptive insulin hypersecretion. These findings suggest that this epigenetic mechanism of nutrient-induced β-cell functional adaptation is relevant during both short-term and chronic nutrient stimulation.

Adaptation of metabolic tissues to changing nutrient states requires accurate interpretation of environmental signals. We show that in β-cells, the chromatin modifier Lsd1 is required for interpreting environmental nutrient signals at the level of the epigenome. The observed enrichment of T2D-associated variants at LSD1-bound sites in human islets suggests that interpretation of nutrient signals is influenced by genetic variation, thereby impacting T2D risk. Gene-environment interactions play an important role in T2D pathogenesis^81^. Our findings support a model whereby influences from environmental nutrient signals and genetic variation converge at the level of the β-cell epigenome to affect adaptation of the insulin secretory response. Deeper investigation of environmental regulation of the epigenome in metabolic tissues should have relevance for understanding the pathogenesis of T2D, and should help pave the way for therapeutic intervention.

## Methods

### Animal studies

All mice used were of mixed strain backgrounds with approximately equal contributions from C57BL/6N and CD1, with the exception of *db*/*db* mice, which were on a pure C57BLKS/J background, and mice used for Lsd1 ChIP-seq, which were on a pure C57BL/6N background. Animals were housed 1-5 mice per cage under standard housing conditions with free access to water and chow (PicoLab Rodent Diet 20 pellets, #5053), unless otherwise indicated. Male mice were used for all experiments unless otherwise indicated. The following mouse strains were used in this study: *Lsd1^flox^*^82^, *Pdx1-CreER*^83^, *MIP-CreER*^84^, *mIns1-H2b-mCherry*^85^, and *db*/*db* (Jackson labs strain 000642). To delete *Lsd1* in *Lsd1^flox/flox^*; *Pdx1-CreER* or *Lsd1^flox/flox^*; *MIP-CreER* mice, tamoxifen (Sigma) was injected subcutaneously with 4 doses of 4 or 6 mg, respectively, every other day in 10- to 14-week-old mice. For *Lsd1* deletion, control *Lsd1^+/+^*; *Pdx1-CreER* or *Lsd1^+/+^*; *MIP-CreER* mice were tamoxifen-injected in parallel unless otherwise indicated. Tamoxifen was dissolved in corn oil at 20 mg/mL for injections. Post-tamoxifen time points are expressed relative to the final tamoxifen injection. *db*/*db* mice were analysed weekly by blood glucose measurements, and islets were harvested from experimental *db/db* and control *db*/+ littermates for ChIP-seq and mRNA-seq once glycaemia in *db/db* mice exceeded 250 mg/dL. All animal experiments were approved by the Institutional Animal Care and Use Committee of the University of California, San Diego.

### Metabolic Studies

Feeding entrainment was adapted from^25^ with the exception that the feeding period encompassed the entire 12-hour dark phase. For all fasting experiments, mice were transferred to fresh cages containing woodchip bedding to prevent coprophagia. Unless otherwise indicated, blood glucose, body weight, and serum insulin were measured between ZT3 and ZT4 (Bayer Contour glucometer; Bayer, Tarreytown, NJ). Food intake was measured by weighing food hoppers over time, whereas each cage of co-housed mice was considered one replicate, and food intake was averaged over the number of mice per cage. Overnight fasts were performed for 16 hours between ZT12 and ZT4 the following day. For serum hormone measurements, blood was collected from the tip of the tail and centrifuged, and the supernatant assayed for insulin content by ELISA (ALPCO) or for glucagon content by radioimmunoassay (EMD Millipore).

Glucose and insulin tolerance tests were performed in mice fasted for 6 hours. Food was withdrawn at ZT0 and fasting blood glucose levels were recorded at ZT6. Mice were then injected intraperitoneally with a dextrose solution at a dose of 1.5 g/kg body weight (glucose tolerance tests) or an insulin solution (bovine insulin, Sigma) at a dose of 0.25 U/kg body weight (insulin tolerance tests). Blood glucose was measured as above at the indicated time points following injections. Pancreatic insulin content was determined as previously described^86^.

### Cell line and islet culture

Mouse islets were isolated and cultured as previously described^45^. Mouse islets were cultured in complete media (RPMI 1640 with 8 mM glucose, 10% FBS, 2 mM L-glutamine, 100 U/mL Pen/Strep, 1 mM sodium pyruvate, 10 mM HEPES, and 0.25 mg/mL amphoterecin B) at 37**°**C with 5% CO_2_ for a maximum of 48 hours. For cAMP stimulation, islets were treated with 0.5 mM cpt-cAMP (Sigma) in culture media containing 11 mM glucose after overnight recovery from isolation. For Erk inhibition, 1 µM SCH772984 (Apex) or vehicle (DMSO) was added to the culture medium for 24 hours immediately following islet isolation and was excluded during perifusion assays.

Human islets were acquired through the Integrated Islet Distribution Program. Upon receipt, islets were stained with dithizone and positively-staining material was hand-picked using a dissection microscope. Islets were allowed to recover 18-48 hours in complete media (CMRL 1066 with 13 mM glucose, 10% FBS, 2 mM L-glutamine, 100 U/mL Pen/Strep, 1 mM sodium pyruvate, 10 mM HEPES, and 0.25 mg/mL amphoterecin B) at 37**°**C with 5% CO_2_ prior to treatments. For LSD1 inhibition studies, 2 mM tranylcypromine (Sigma) or vehicle (H_2_O) was included in the culture media for 24 hours prior to analysis. Donor information can be found in Supplementary Table 9.

### Immunohistochemistry, islet composition, and β-cell mass measurements

Immunohistochemistry, islet endocrine cell composition, and β-cell mass measurements were performed as previously described^86^. A mouse-on-mouse kit (Vector labs) was used according to the manufacturer’s instructions to stain using primary antibodies of mouse origin.

### Insulin secretion measurements

Static insulin secretion assays were performed as previously described^45^. Unless otherwise noted, all insulin secretion measurements were performed after islets were allowed to recover overnight from isolation (mouse islets) or shipment (human islets) in complete islet media (formulations described above). Secreted insulin and islet insulin content were determined by ELISA (ALPCO) according to the manufacturer’s instructions.

Insulin secretion assays were performed immediately following islet isolations and hand-picking for comparison of islets in the fed and fasted states and for comparison of *db/db* and control (*db/+*) islets to avoid reversion of transient in vivo changes. For comparison of islets in the fed and fasted states, fatty-acid-free bovine serum albumin was used to avoid the stimulatory effect of trace fatty acids in the fasted state^27^.

The following compounds were included as indicated during stimulations: Exendin-4 (10 nM), L-leucine and L-glutamine (10 mM each), the fatty acid palmitate (100 µM), KCl (30 mM), and diazoxide (100 µM). For glycolysis inhibition, islets were preincubated in 100 mM D-mannoheptulose (Fisher Scientific) in Krebs-Ringers-Bicarbonate-HEPES (KRBH) buffer (130 mM NaCl, 5 mM KCl, 1.2 mM CaCl_2_, 1.2 mM MgCl_2_, 1.2 mM KH_2_PO_4_, 20 mM HEPES pH 7.4, 25 mM NaHCO_3_, and 0.1% bovine serum albumin) containing 2.8 mM glucose for one hour at 37**°**C with 5% CO_2_ prior to the beginning of the experiment and where indicated, mannoheptulose continued to be included throughout starvation and stimulation steps.

Perifusion was carried out using the Biorep perifusion system. Islets were first starved for 30 min in KRBH containing 2.8 mM glucose at 37**°**C with 5% CO_2_ and were then loaded into perifusion chambers (130 islets/chamber). For equilibration, islets were perifused with KRBH containing 2.8 mM glucose for 30 min, at which point islets were stimulated with KRBH containing the indicated glucose concentrations and perifusate was collected for analysis. Perifusion was carried out at 37**°**C with a flow rate of 80 µL/min. At the end of each experiment, islets were transferred to Eppendorf tubes and lysed by sonication for insulin content determination.

### Islet respirometry

Respirometry was performed as previously described^45^. Briefly, following islet isolations, islets were pooled within each experimental group and transferred to complete islet media, then shipped overnight at room temperature to Boston University. Islets were then allowed to recover overnight at 37**°**C with 5% CO_2_, then respirometry was performed using the Seahorse Bioscience XF24 Extracellular Flux Analyzer. Plates were first loaded with 60 islets per well in XF Assay Media (Seahorse Bioscience) supplemented with 2.8 mM glucose, 1% FBS, and 5 mM HEPES. Following a 1-hour starvation period in this media, oxygen consumption was measured at baseline and following sequential additions of 16.8 mM glucose, 5 µM oligomycin, and 2 µM antimycin A.

### ATP measurements

Following islet isolations, islets were pooled within each experimental group then allowed to recover overnight in complete media. The day of the assay, islets were pre-starved in KRBH containing 2.8 mM glucose for 1 hour, and then transferred to 200 µl of the same media as pools of 20 islets each in round-bottom 96 well plates and incubated for 30 min to allow for ATP production. Islets were then washed once in ice cold DPBS then lysed in 0.83% trichloroacetic acid (TCA) for 5 min at room temperature. Following lysis, TCA was neutralized with 60 mM Tris-Acetate pH 8.0 and 0.6% SDS. ATP measurements were performed using the Enliten ATP assay kit (Promega) according to the manufacturer’s instructions.

### Calcium imaging

Calcium imaging was performed with the indicated genotypes of mice also carrying the *mIns1-H2b-mCherry* transgene^85^. Following islet isolations, islets were allowed to recover overnight in complete islet media at 37**°**C with 5% CO_2_. Islets were dispersed with 0.05% trypsin/EDTA (Invitrogen) by incubating at 37**°**C for 5 min with gentle agitation, then dissociated islet cells were plated on glass-bottom petri dishes coated with poly-l-lysine. Cells were allowed to attach overnight. The day of the assay, cells were loaded with Fluo-4 by incubation in KRBH containing 5.5 mM glucose, 4.56 µM Fluo-4, AM (Thermo), and 0.02% Pluronic F-127 (Biotium) for 45 min at 37**°**C. Cells were then washed three times with KRBH containing 2.8 mM glucose and were starved for 1 hour prior to imaging. Fluorescence was monitored and analysed as previously described^87^. Fluorescence maxima and minima were obtained by treating cells with 50 µM ionomycin or 100 mM EGTA, respectively.

### Targeted metabolomics

Islets were prepared for targeted metabolomics essentially as previously described^45^. Briefly, islets were pooled within each experimental group and allowed to recover overnight in complete islet media. The day of nutrient tracing, islets were briefly washed in KRBH containing 2.8 mM unlabeled glucose and were then transferred to KRBH containing 2.8 mM U-^13^C glucose in low-adherence 12-well plates in groups of 220 islets per well (each well being considered a replicate). Plates were then incubated at 37**°**C and 5% CO_2_ for two hours to allow for ^13^C incorporation into downstream metabolites, which is not expected to result in steady state labelling based previous islet tracing experiments^45^. After tracing, islets were transferred to pre-chilled Eppendorf tubes, washed once in 0.9% NaCl, and snap frozen. Samples were prepared for GC-MS and analysed as previously described^45^. The same numbers of islets were extracted in each group and norvaline was added as an internal standard. Data was normalized to the internal standard and metabolite abundances were expressed relative to control islets.

### Human β-cell dissociation/reaggregation, lentiviral shRNA transduction, and insulin secretion assays

Immediately following shipment, human islets were dissociated by first washing in Versene (Thermo) and then incubating with 0.05% trypsin/EDTA at 37 **°**C for 7 minutes. Islet pellets were triturated then digestion was quenched in islet media. Dissociated islet cells were pelleted via centrifugation for 5 minutes at 1200 rpm in a swing-out rotor centrifuge, and then resuspended in CMRL media supplemented with 5% foetal bovine serum, 20 mM HEPES, 1 mM EDTA, and 0.1 mg/mL DNase. Islet cell suspensions were stained with HIC3-2D12 (1:20 dilution) and APC-conjugated HIC1-2B4 (1:100) antibodies. HIC-2D12 was detected with anti-IgM PE. Cells were then stained with propidium iodide as a dead cell marker, and live β-cells were FACS-sorted based on negativity for propidium iodide and HIC-2D12 and positivity for HIC1-2B4, as described elsewhere^88^. Collected β-cells were immediately pelleted by centrifugation as before, then resuspended at a concentration of 50,000 cells/mL. pLKO.1-encoded lentiviruses producing either nontargeting scramble shRNA or a pool of lentiviruses producing *LSD1* shRNA were then added to cells. *LSD1* shRNA mature antisense sequences are as follows: Nontargeting control, CCTAAGGTTAAGTCGCCCTCG; TRCN0000046068, TATTCAGTTTAATGTCTAGGC; TRCN0000046069, TAGTGCCAACAGTATTGGAGC; TRCN0000046070, TAAGGTGCTTCTAATTGTTGG; TRCN0000046071, ATGACTAAGGTAAGATGTAGC.

Cells were seeded at 5,000 cells/well into V-bottom 96-well plates. Plates were spun at 365 rcf for 5 minutes at room temperature to pellet islet cells, and were then incubated overnight at 37**°**C with 5% CO_2_. Islets were then subjected to a complete media change and incubated an additional 4 days to allow for β-cell reaggregation and LSD1 protein degradation, at which point insulin secretion assays were performed. For insulin secretion assays, islet aggregates were washed three times with KRBH supplemented with 2.8 mM glucose then incubated at 37**°**C with 5% CO_2_ for one hour for starvation, at which point starvation media was removed. Aggregates were then incubated with KRBH containing either 2.8 mM glucose or 16.8 mM glucose and returned to the incubator for one hour. Following incubation, media and lysates were harvested as described above for aggregates and insulin content was determined by ELISA (ALPCO) according to the manufacturer’s instructions. Knockdown was verified in whole dissociated islets infected as above and reaggregated in Aggrewell plates, using GFP-expressing pLKO.1-encoded lentivirus for visualization of transduction efficiency in control islets. Reaggregated whole islets were visualized for GFP expression and harvested for Western blotting 72 hours following transduction.

### RNA extraction, mRNA-seq, and qRT-PCR

Unless otherwise stated, islets were processed for RNA purification immediately following isolation. Islets were hand-picked twice under a dissecting microscope to minimize acinar contamination. Islets were then pooled from at least 2 mice and RNA was isolated using the RNeasy Micro kit (QIAGEN) according to the manufacturer’s instructions. mRNA-seq libraries were prepared from 35 ng of total RNA using the TruSeq Stranded mRNA Library Prep Kit (Illumina) with the exception of *db/db* mouse islets and their respective controls, from which libraries were generated with 1 µg of total RNA as previously described^89^. Libraries were single-end sequenced at a length of 50 bp per read using HiSeq 4000 (Illumina). RT-qPCR was performed as previously described^86^. A list of primers used for qPCR can be found in Supplementary Table 10.

### ChIP-seq

Islets were processed for ChIP immediately following isolation (for mouse islets) or shipping (for human islets). Mouse islets were hand-picked twice under a dissecting microscope to minimize acinar cell contamination. Human islets selected for ChIP-seq analysis were of ≥ 90% purity to facilitate analysis of large quantities of islets without the need for dithizone staining. For Lsd1 ChIP, islets were transferred to complete islet media containing 1.11% formaldehyde and fixed on a rocker for 15 minutes. The reaction was quenched for 5 minutes in 0.125 M glycine. Cells were then washed in DPBS containing 0.5% NP-40 then once again in DPBS supplemented with 0.5% NP-40 and 1 mM PMSF. Islets were lysed and ChIP was performed using the ChIP-IT High Sensitivity Kit (Active Motif) with 30 µg of sheared chromatin and 4 µg anti-Lsd1 antibody (ab17721, Abcam). For histone ChIP, islets were washed once in Hanks Balanced Salt Solution (Hyclone) and then fixed for 10 min in DPBS containing 1.11% formaldehyde. Cross-linking was quenched as before, then islets were lysed by passage through a 25 gauge needle in lysis buffer (10 mM Tris-Hcl, pH 8.0, 10 mM NaCl, 3 mM MgCl_2_, 1% NP-40, 1% SDS, 0.5% sodium deoxycholate). ChIP was then performed as described above using 10 or 30 µg of sheared chromatin and 4 µg of anti-H3K4me1 (ab8895, Abcam) or anti-H3K27ac (39133, Active Motif) antibodies, respectively. Libraries were constructed from purified DNA using the KAPA DNA Library Preparation Kit for Illumina (Kapa Biosystems). Input libraries were prepared from each experimental replicate using 10 ng of DNA purified immediately following shearing. Libraries were sequenced using HiSeq 4000 (Illumina) as above.

### mRNA-seq data analysis

mRNA-seq reads were mapped to NCBI37/mm9 (mouse) or GRCh37/hg19 (human) genomes by STAR (STAR-STAR_2.4.0f1, --outSAMstrandField intronMotif -- outFilterMultimapNmax 1 --runThreadN 5), excluding reads mapping to multiple loci. Reads with exact matches were used to determine RPKM by Cufflinks (cufflinks v2.2.1,- p 6 -G $gtf_file --max-bundle-frags 1000000000). Correlations between mRNA-seq replicates can be found in Supplementary Table 1. Genes with mean RPKM ≥ 1 in at least one experimental group were considered to be expressed, and all non-expressed genes were excluded from downstream analyses. According to this criterion, 12,447 genes were expressed in fed or fasted islets, 12,763 genes were expressed in Lsd1^Δβ^ or control islets over the combined time points, and 12,750 genes were expressed in *db/db* or control (*db/+*) islets. Cuffdiff was used to assess expression differences for all pairwise comparisons, with *P* < 0.01 considered significant. For volcano plots, *P*-values = 0 were graphed as 0.00001. GSEA was performed using default settings with an FDR cutoff of 0.25 being considered significant^90^. Enrichment in human gene sets was performed by converting human gene symbols to mouse orthologues using bioDBnet^91^. All differentially expressed genes identified in this study are listed in Supplementary Table 2a-d.

### ChIP-seq data analysis

Single-end 50-bp ChIP-seq reads were mapped to NCBI37/mm9 (mouse) or GRCh37/hg19 (human) genomes using Bowtie2 with a seed length of 33 bp and a maximum of 2 mismatches allowed in the seed region, discarding reads aligning to multiple sites. Duplicate reads were removed using SAMtools. Biological replicates from each condition (*n* = 2) were assessed for similarity genome-wide using the multiBamSummary program (in “bins” mode) of the deepTools bioinformatics suite^92^. Replicates were merged for analysis based on high (≥ 0.78) Pearson correlation coefficients as indicated in Supplementary Table 3. Human islet H3K27ac ChIP-seq data were downloaded from^93,94^ and processed in parallel.

Tag directories were generated from bam files and replicates were combined using HOMER. ChIP-seq peaks within tag directories were identified with the findPeaks command in HOMER (-style histone, -P 0.0000001 for H3K27ac and -style factor for Lsd1) using input tag directories as background. For identification of changes in H3K27ac signal with feeding, all H3K27ac peaks in the fed and fasted states were merged using BEDtools and then findPeaks was again used to identify peaks with higher signal (≥ 1.2-fold and *P*-value < 0.0001) in fed or fasted H3K27ac tag directories with the reciprocal condition set as background. The annotatePeaks.pl command in HOMER was used to quantify tag densities for histograms, heatmaps, and boxplots. Tag density boxplots (± 1 kb from the center of each peak) were visualized using R, with boxplot whiskers indicating the interquartile range x 1.5 or maxima and minima if all data points fell within this range. Heatmaps were visualized using the heatmap.2 package in R. BEDtools was used to determine associations between different coordinate sets (e.g. Lsd1 or H3K27ac peaks) located within ± 1 kb. ChIP-seq genome browser tracks were generated using the makeUCSCfile command in HOMER, which normalizes all tag directories to 10^7^ total tags. A smoothening window of “4” was applied to all tracks using the UCSC Genome Browser.

Classes of Lsd1-regulated H3K27ac peaks were determined based on tag intensity calls ± 1 kb using the annotatePeaks.pl command in HOMER as described for boxplots above. To normalize distribution, each tag density value was transformed using the formula log_2_((y + 1)/(x + 1)), with x representing the control tag density value and y representing the Lsd1^Δβ^ tag intensity value. Peaks were then categorized based on the time point at which they first reached the cutoff of 1.5-absolute fold difference in either direction, resulting in classes of peaks that gain H3K27ac first at week 1 (3,320 peaks), peaks that gain H3K27ac first at week 3 (10,233), peaks that lose H3K27ac first at week 3 (2,713), peaks that never reach this cutoff (21,972), and a negligible number of peaks that lose H3K27ac first at week 1 (44) or exhibit losses and then gains in H3K27ac at week 1 and week 3, respectively (14). Peaks that first gain H3K27ac at week 1 were then further subdivided into peaks that continue to accumulate H3K27ac from week 1 to week 3 (≥ 0.5-absolute fold additional increases over week 1; 448 peaks), peaks that exhibit concordant gains from Wk 1 to Wk 3 (within ± 0.5-absolute fold additional changes; 2,611 peaks), and into a negligible number of peaks that gained then lost H3K27ac from week 1 to week 3 (261 peaks). Further analysis excluded classes represented by negligible numbers of peaks as indicated above, focusing on those described in Extended Data Fig. 5a.

Classes of Lsd1-regulated H3K4me1 patterns at active chromatin were determined similarly to those of H3K27ac. Considering that H3K4me1 exhibited more modest changes relative to H3K27ac, thresholds were set to 1.2-absolute fold differences. H3K4me1 changes were more concordant from week 1 to week 3 and therefore all time points and directions of deregulation were considered: peaks gaining H3K4me1 at both time points (9,780), gaining H3K4me1 at week 1 only (9,988), gaining H3K4me1 at week 3 only (2,569), losing H3K4me1 at both time points (1,640), losing H3K4me1 and week 1 only (1,756), losing H3K4me1 at week 3 only (1,709), or failing to reach the threshold at any time point (10,283). Coordinates of all ChIP-seq peaks and peak classes can be found in Supplementary Tables 4a-d and 6a-c.

Heatmap data of relative H3K4me1 in Lsd1^Δβ^ islets were generated by transforming 100 bp bin tag densities using the formula log_2_((y + 0.05)/(x+0.05)) to match transformations applied over a 2 kb window for the above analyses. Heatmap data was sorted based on Lsd1 tag counts and displayed using the heatmap.2 package in R.

### Motif enrichment analysis

Motif enrichments were determined using HOMER. Enrichments were calculated for sequences within peak coordinates (-size given) relative to the background set of H3K27ac peaks indicated for each calculation. A complete list of motif enrichment results can be found in the Supplementary Table 5a-b.

### Permutation-based significance

Enrichment tests for associations among TSSs, Lsd1 peaks, and H3K27ac peaks were determined using a random sampling approach to compare the number of true overlaps to the number of expected overlaps. Null distributions (expected overlap frequencies) were obtained by performing 10,000 iterations of randomly shuffling test coordinates using BEDtools then intersecting shuffled coordinates with reference coordinates ± 50 kb. Permutation data plots are presented with test coordinate sets on the x-axis and reference coordinate sets as the plot title.

### Gene ontology

Functional categories related to the set of feeding-regulated genes and links between each pair of categories were identified with Metascape as described in http://metascape.org/. Statistically enriched pathways from KEGG and Reactome were hierarchically clustered into a tree based on Kappa-statistical similarities among their gene memberships. A 0.3 kappa score was applied as the threshold to define clusters, each of them representing a group of similar functional categories. A subset of representative terms from each cluster was automatically selected by Metascape and converted into a network, where terms with similarity > 0.3 are connected by edges. Specifically, terms with the most significant *P*-values from each of the clusters were depicted as network nodes, with the constraint of having a maximum of 15 terms per cluster and 250 terms in total, with node size representative of the degree of enrichment. For clarity, a representative term was selected to represent each cluster.

### Genome-wide association studies (GWAS) enrichment methods

Stratified LD score regression^95^ was used to assess whether LSD1-bound sites were enriched for GWAS signal for metabolic (HOMA-B^64^, HOMA-IR^64^, fasting glucose^65^, fasting glucose adjusted for BMI^65^, fasting insulin^65^, fasting insulin adjusted for BMI^65^, 2 hour glucose adjusted for BMI^66^, fasting proinsulin^67^, and type 2 diabetes^63^) and well-powered non-metabolic control traits (Alzheimer’s disease^68^, systemic lupus erythematosus^69^, autoimmune vitiligo^70^, primary biliary cirrhosis^71^, primary sclerosing cholangitis^72^, inflammatory bowel disease^73^, rheumatoid arthritis^74^, coronary artery disease^75^, schizophrenia^76^, and major depressive disorder^77^), using European subset summary statistics where available. After filtering out ENCODE blacklisted regions^34^, islet LSD1 peaks were used as a binary annotation. The partitioned heritability version of LD score regression was used to estimate enrichment (%h^2^/%SNPs) using the baseline LD model v2.2.

Putative target genes of LSD1-bound sites were determined using promoter capture Hi-C data from primary human islets^23^. For each site, gene promoters mapping in a chromatin loop to the site were identified using a 5 kb flanking window around loop anchors. Genetic variants in T2D and fasting glucose adjusted for BMI GWAS datasets were intersected with LSD1-bound sites, retaining variants with at least nominal association (*P* < 0.05), and annotating variants with putative target genes of the overlapping LSD1 site.

## Acknowledgements

We thank A. Pospisilik and S. Panda for helpful discussions, N. Rosenblatt for mouse husbandry, I. Matta for assistance with islet preparations, and Michael Rosenfeld for *Lsd1^flox^* mice. We acknowledge support of the UCSD IGM Genomic Center (P30 DK064391) for mRNA-seq and ChIP-seq. Human pancreatic islets were provided by the NIDDK-funded Integrated Islet Distribution Program (IIDP) at City of Hope, National Institutes of Health (NIH) Grant 2UC4DK098085. This work was supported by NIH training grant T32DK007494-30, Juvenile Diabetes Research Foundation postdoctoral fellowship 3-PDF-2014-193-A-N, and John G. Davies Endowed Fellowship in Pancreatic Research to M. Wortham, NIH grants DK068471 to M.S., CA188652 to C.M.M., DK114650 to K.J.G., DK110276 to M.O.H., and DK116038 to U.S.J.

## Author Contributions

M. Wortham and M.S. conceived the study, were responsible for its overall design, and prepared the manuscript. M. Wortham, F.L., and J.Y.F. performed mouse experiments. F.L. performed islet isolations. M. Wortham, J.Y.F., A.R.H., B.R.C., N.A.P., and M.O.H. performed islet experiments and hormone measurements. M. Wortham, N.K.V., N.A.P., and U.S.J. generated ChIP-seq data. M. Wortham, M. Wallace, and C.M.M. performed and/or interpreted glucose tracing experiments. M. Wortham, F.M., N.K.V., and Y.S. performed bioinformatics analysis. J.C. and K.J.G. analysed human genetic data. O.S.S. designed and interpreted islet respirometry experiments.

## Author Information

The authors declare no competing interests. Correspondence and requests for materials should be addressed to.

## Data availability

ChIP-seq and mRNA-seq data will be deposited in GEO prior to publication. Accession numbers for human islet H3K27ac ChIP-seq data used in this study are: GSE51311 and E-MTAB-1919.

## Extended Data

**Extended Data Figure 1.**
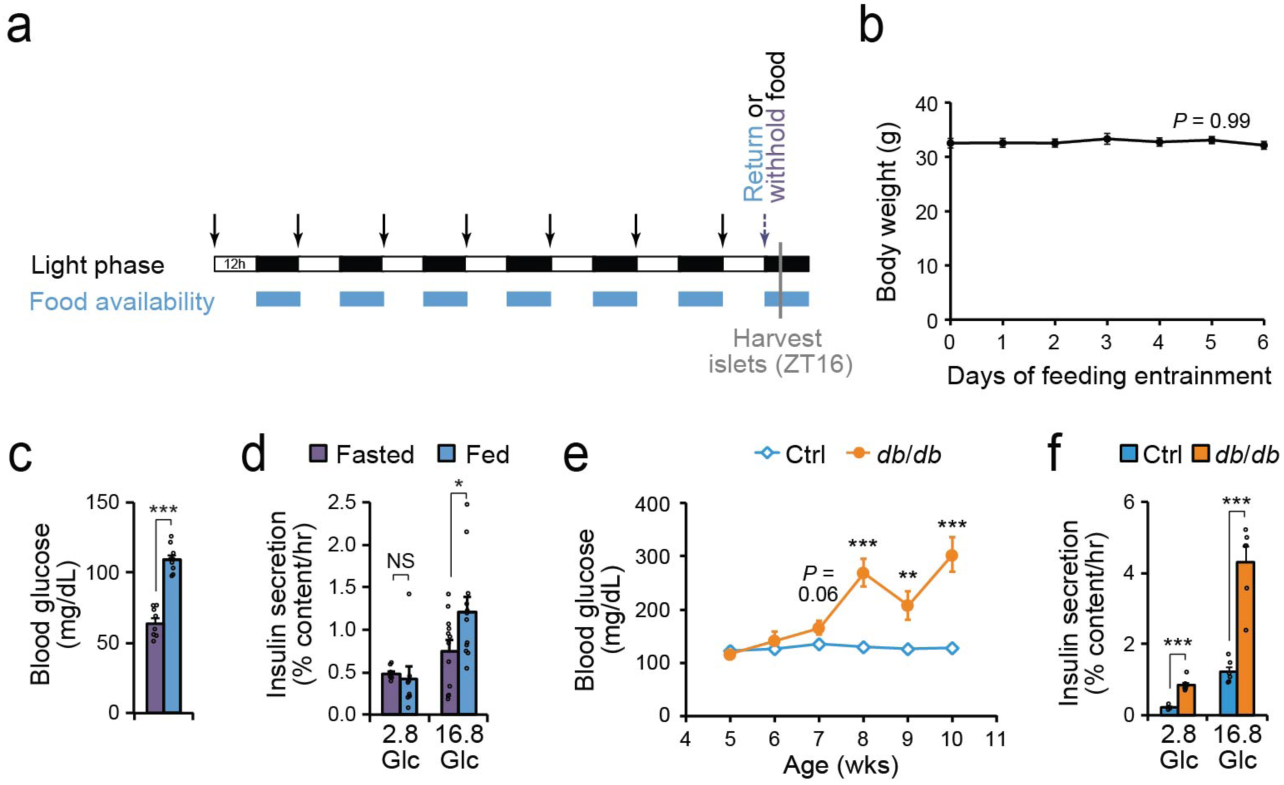
Short-term and prolonged increases in nutrient intake enhance glucose stimulated insulin secretion. Related to Figure 1. **(a)** Schematic of time-restricted feeding experiment used to entrain food intake in mice. Arrows indicate food removal and body weight measurements. ZT, Zeitgeber time; h, hours. **(b)** Animal weights at the indicated days of time-restricted feeding (taken at ZT0). *n* = 20 mice. *P*-value was calculated by one-way ANOVA. **(c)** Blood glucose levels at the time of islet isolation (ZT16). Fasted: *n* = 8 mice, fed: *n* = 9 mice. \*\*\**P* < 0.001 by unpaired two-tailed t-test. **(d)** Static insulin secretion assay in islets from fed and fasted mice stimulated with the indicated glucose (Glc) concentrations (in mM). Fasted or fed 2.8 mM glucose: *n* = 8 pools of 10 islets each, fasted or fed 16.8 mM glucose: *n* = 12 islet pools. \**P* < 0.05, by unpaired two-tailed t-test with Welch’s correction for unequal variance as necessary. **(e)** Time course of ad libitum-fed blood glucose levels in control (ctrl, *db*/+; *n* = 19) and *db*/*db* (*n* = 18) mice at the indicated ages in weeks (wks). Data are shown as mean ± S.E.M. \*\**P* < 0.01, \*\*\**P* < 0.001 by unpaired two-tailed t-test with Welch’s correction for unequal variance as necessary. **(f)** Static insulin secretion assay in 10-week-old control and *db*/*db* islets stimulated with the indicated glucose (Glc) concentrations (in mM). Control 2.8 mM glucose: *n* = 4 pools of 10 islets each, all other groups: *n* = 6 islet pools. Data in are shown as mean ± S.E.M. \*\*\**P* < 0.001 by unpaired two-tailed t-test with Welch’s correction for unequal variance. Data are shown as mean ± S.E.M. Source data for all quantifications and exact *P*-values for all indicated statistical tests are provided online.

**Extended Data Figure 2.**
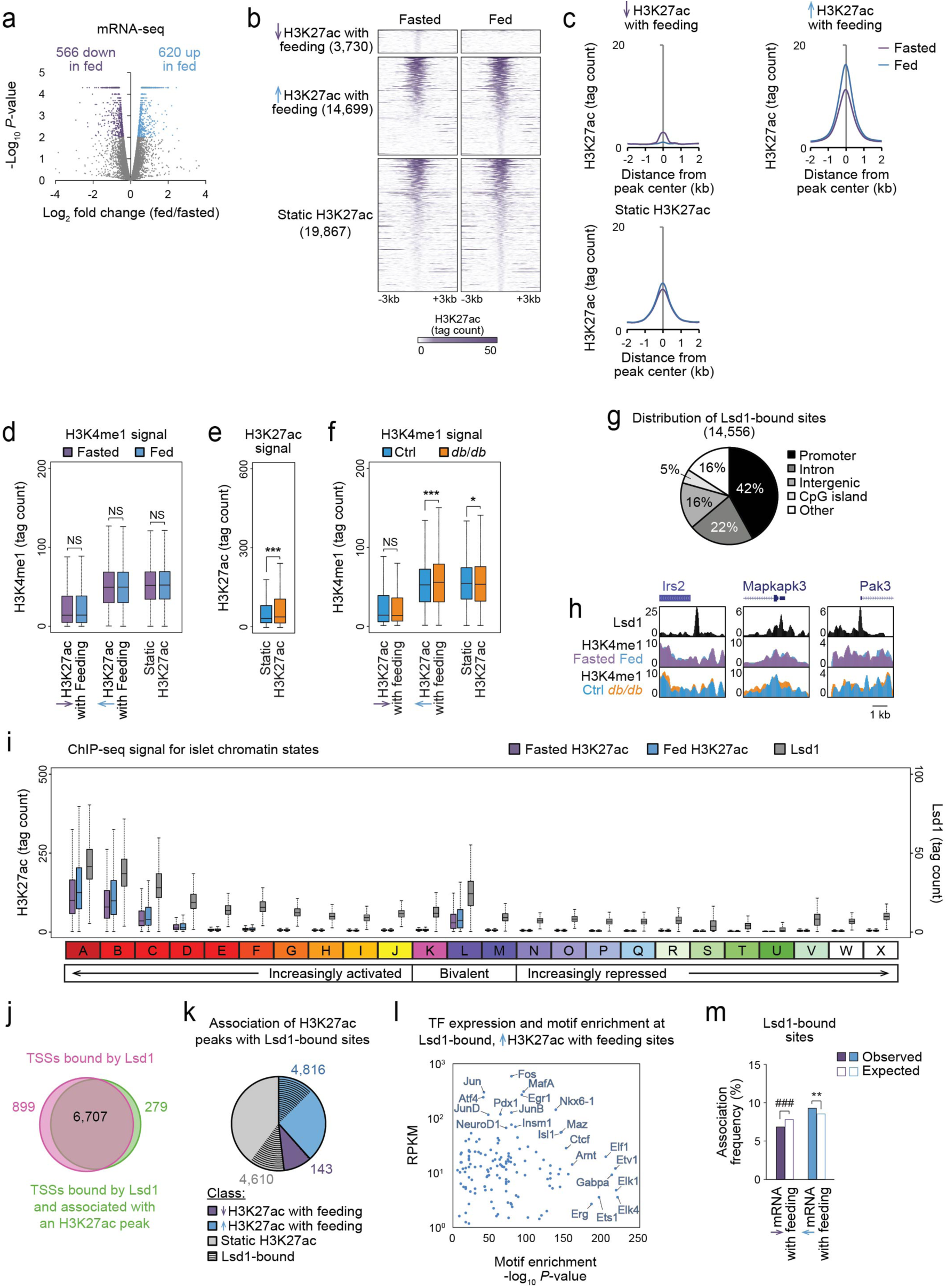
Lsd1 associates with feeding-regulated active chromatin enriched for motifs recognized by signal-dependent transcription factors. Related to Figure 1. **(a)** Volcano plot comparing mRNA levels in islets from fed and fasted mice. Differentially expressed genes are indicated in purple or blue (*P* < 0.01 by Cuffdiff). *n* = 3 biological replicates of islets pooled from separate mice per group. **(b** and **c)** Heatmaps (b) and histograms (c) of H3K27ac ChIP-seq signal for the indicated classes of H3K27ac peaks. *n* = 2 biological replicates of islets pooled from separate mice per group; data from independent ChIP-seq experiments were merged for analysis. **(d-f)** ChIP-seq signal for H3K4me1 (d, f) and H3K27ac (e) at the indicated classes of H3K27ac peaks in islets from fed and fasted mice (d) or in *db/db* and control (ctrl, *db*/+) islets (e, f). *n* = 2 biological replicates of islets pooled from separate mice per group; data from independent ChIP-seq experiments were subsequently merged for analysis. Boxplot whiskers span data points within the interquartile range x 1.5. \**P* < 0.05, \*\*\**P* < 0.001 by Wilcoxon rank-sum test; NS, not significant. **(g)** Distribution of Lsd1 peaks at the indicated genomic features. Lsd1 ChIP-seq data are from *n* = 1 biological replicate from pooled islets. Data were highly correlated with Lsd1 ChIP-seq data from an independent biological replicate (see Supplementary Table 3). **(h)** Lsd1 and H3K4me1 ChIP-seq genome browser tracks for genes shown in Fig. 1o in islets from fed and fasted mice and in *db/db* and control islets. **(i)** ChIP-seq signal for H3K27ac and Lsd1 at the indicated chromatin states. **(j)** Venn diagram comparing the number of transcription start sites (TSSs) associated ± 50 kb with an Lsd1 peak and TSSs associated ± 50 kb with an H3K27ac peak within ± 1 kb of an Lsd1 peak. **(k)** Proportion of the indicated classes of H3K27ac peaks associating with an Lsd1 peak ± 1 kb. Numbers indicate H3K27ac peaks associated with an Lsd1 peak. **(l)** Transcription factor (TF) motifs enriched at H3K27ac peaks gaining acetylation with feeding and associated with an Lsd1 peak ± 1 kb relative to H3K27ac peaks gaining acetylation with feeding and not associated with an Lsd1 peak plotted against mRNA levels of cognate TFs in islets from fed mice. **(m)** Association frequencies between TSSs of genes downregulated or upregulated by feeding with an Lsd1 peak ± 50 kb. Open bars indicate association frequencies expected by chance. \*\**P* < 0.01 for enrichment and ^###^P < 0.001 for depletion by permutation test. Source data for all quantifications and exact *P*-values for all indicated statistical tests are provided online.

**Extended Data Figure 3.**
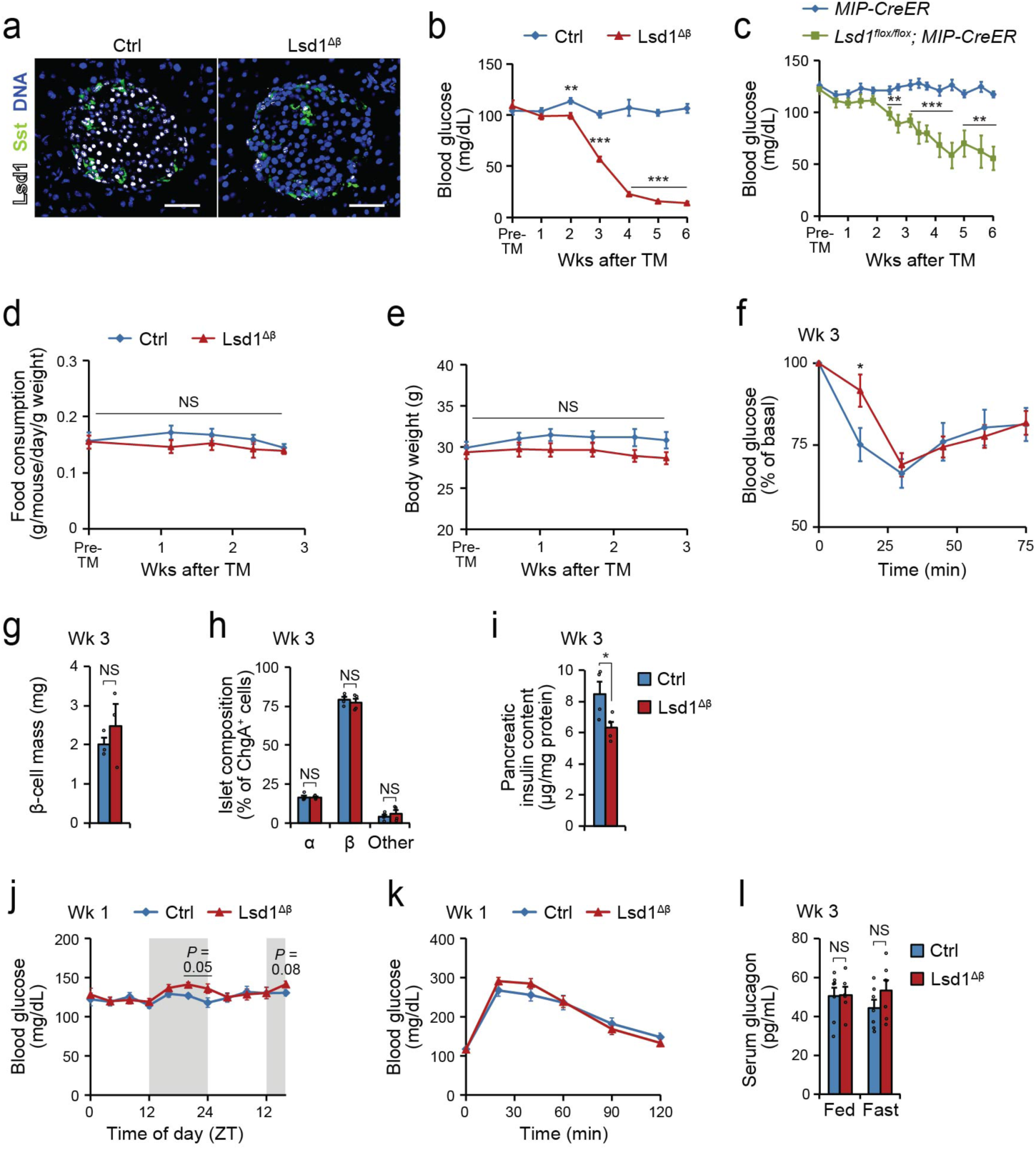
Hypoglycaemia in Lsd1^Δβ^ mice is not caused by altered insulin sensitivity, food intake, β-cell mass, or circulating glucagon. Related to Figure 2. **(a)** Immunofluorescence staining for the indicated proteins on pancreas sections from control (ctrl, TM-treated *Lsd1^flox/+^*; *Pdx1-CreER* mice) and Lsd1^Δβ^ mice two days after TM treatment. Scale bar, 50 µm. **(b)** Time course of ad libitum-fed blood glucose levels in female control (TM-treated *Lsd1^+/+^*; *Pdx1-CreER* mice, *n* = 11) and Lsd1^Δβ^ (*n* = 10) mice. pre-TM, within 3 days prior to initial TM injection. **(c)** Time course of ad libitum-fed blood glucose levels in *Lsd1^+/+^*; *MIP-CreER* (*n* = 12) and *Lsd1^flox/flox^; MIP-CreER* (*n* = 8) mice following TM treatment. **(d** and **e)** Time courses of food intake (d) and body weight (e) in control and Lsd1^Δβ^ mice following TM treatment. *n* = 5 cages of mice per group (d), *n* = 10 mice per group (e). **(f)** Insulin tolerance tests in control (*n* = 7) and Lsd1^Δβ^ (*n* = 9) mice three weeks following TM treatment. **(g**-**i)** β-cell mass (g), islet cell type composition (h), and pancreatic insulin content (i) of control and Lsd1^Δβ^ mice three weeks following TM treatment. *n* = 3 (g) or 4 (h, i) mice per group. **(j)** Blood glucose levels in control and Lsd1^Δβ^ mice at 4-hour increments across a 40-hour time course in control and Lsd1^Δβ^ mice one week following TM treatment. *n* = 7 mice per group. **(k)** Glucose tolerance tests in control (*n* = 12) and Lsd1^Δβ^ (*n* = 15) mice one week following TM treatment. **(l)** Serum glucagon levels in ad libitum-fed and 16-hour fasted mice of the indicated genotypes. Fed Lsd1^Δβ^: *n* = 6, all other groups: *n* = 7 mice per group. Wk/s, week/s; TM, tamoxifen. Data are shown as mean ± S.E.M. \**P* < 0.05, \*\**P* < 0.01, \*\*\**P* < 0.001 by unpaired two-tailed t-test with Welch’s correction for unequal variance as necessary (b-l). NS, not significant. Source data for all quantifications and exact *P*-values for all indicated statistical tests are provided online.

**Extended Data Figure 4.**
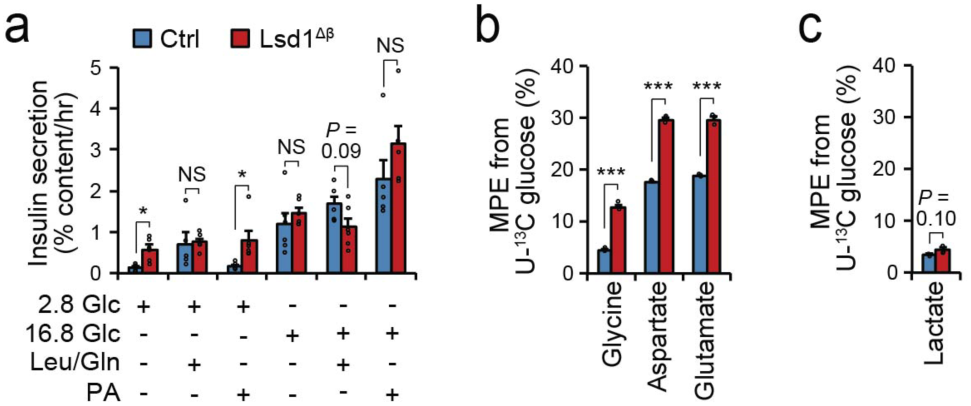
Functional and metabolic characteristics of islets from Lsd1^Δβ^ mice. Related to Figures 2 and 3. **(a)** Static insulin secretion assays in control (ctrl) and Lsd1^Δβ^ islets stimulated with the indicated glucose (Glc) concentrations (in mM) with or without amino acids leucine and glutamine (Leu/Gln) or the fatty acid palmitate (PA). Control 2.8 mM Glucose + Leu/Gln, Control 2.8 mM Glucose + PA, Control 16.8 mM Glucose + Leu/Gln, Control 16.8 mM Glucose + PA, Lsd1^Δβ^ 16.8 mM Glucose + PA: *n* = 5 pools of 10 islets each per group, all other groups: *n* = 6 islet pools. **(b** and **c)** Molar percent enrichment (MPE) of ^13^C for amino acids (b) and lactate (c) in islets following two hours of tracing in 2.8 mM U-^13^C glucose. *n* = 3 pools of 220 islets each. Islets were isolated from control and Lsd1^Δβ^ animals three weeks following tamoxifen treatment for all experiments. Data are shown as mean ± S.E.M. \**P* < 0.05, \*\*\**P* < 0.001 by unpaired two-tailed t-test with Welch’s correction for unequal variance as necessary. NS, not significant. Source data for all quantifications and exact *P*-values for all indicated statistical tests are provided online.

**Extended Data Figure 5.**
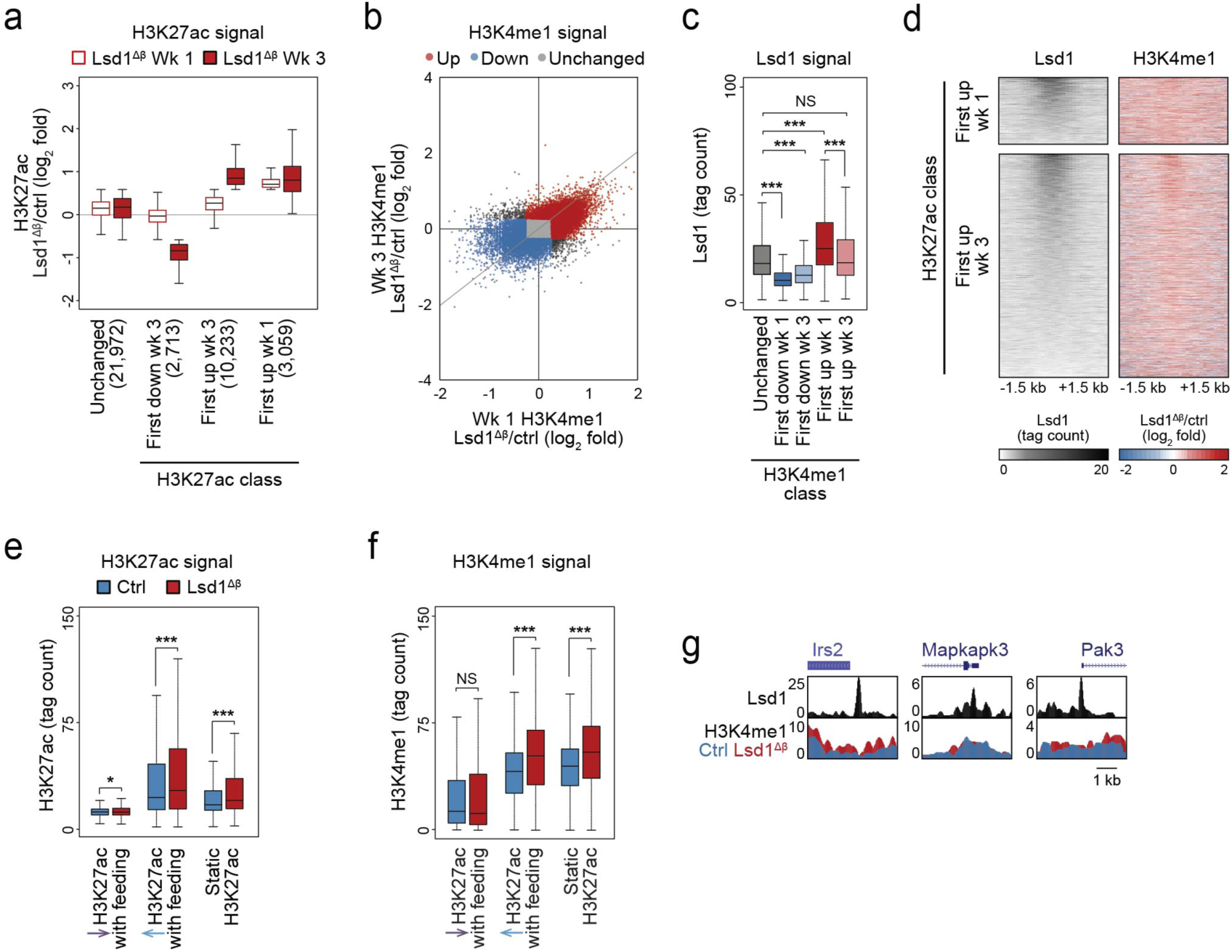
Concordant changes in H3K27ac and H3K4me1 after *Lsd1* inactivation. Related to Figure 4. **(a)** H3K27ac ChIP-seq signal at the indicated classes of H3K27ac peaks (from Fig. 4c) plotted as relative tag density in Lsd1^Δβ^ islets compared to control (ctrl) islets at one and three weeks (Wk) following TM treatment. *n* = 2 biological replicates of islets pooled from separate mice per group; data from independent ChIP-seq experiments were subsequently merged for analysis. Boxplot whiskers span data points within the interquartile range x 1.5. **(b)** H3K4me1 ChIP-seq signal at H3K27ac peaks (from Extended Data Fig. 1b) plotted as relative tag density in Lsd1^Δβ^ islets compared to control islets at the indicated time points following TM treatment. The gray line indicating a slope of 1 provides a reference for corresponding changes at the two time points. Peaks were classified by the timepoint when relative tag density in Lsd1^Δβ^ islets exceeded 1.2-absolute fold difference compared to control islets (see Methods for additional details on peak classification). *n* = 2 biological replicates of islets pooled from separate mice per group; data from independent ChIP-seq experiments were subsequently merged for analysis. **(c)** Lsd1 ChIP-seq signal at the indicated classes of H3K4me1 sites (from b). Lsd1 ChIP-seq data are from *n* = 1 biological replicate from pooled islets. Data were validated to be highly correlated with ChIP-seq data prepared from an independent biological replicate. Boxplot whiskers span data points within the interquartile range x 1.5. **(d)** Heatmaps of Lsd1 ChIP-seq signal and relative H3K4me1 tag density in Lsd1^Δβ^ islets compared to control islets one week following TM treatment for the indicated classes of H3K27ac peaks (from Fig. 4c). Coordinates indicate position relative to each H3K27ac peak center. Peaks are ordered by Lsd1 intensity. **(e** and **f)** ChIP-seq signal for H3K27ac (e) and H3K4me1 (f) at the indicated classes of H3K27ac peaks in control and Lsd1^Δβ^ islets one week following TM treatment. Boxplot whiskers span data points within the interquartile range x 1.5. **(g)** Lsd1 and H3K4me1 ChIP-seq genome browser tracks for genes shown in Fig. 4g in control and Lsd1^Δβ^ islets one week following TM treatment. \**P* < 0.05, \*\*\**P* < 0.001 by Wilcoxon rank-sum test. NS, not significant. Source data for all quantifications and exact *P*-values for all indicated statistical tests are provided online.

**Extended Data Figure 6.**
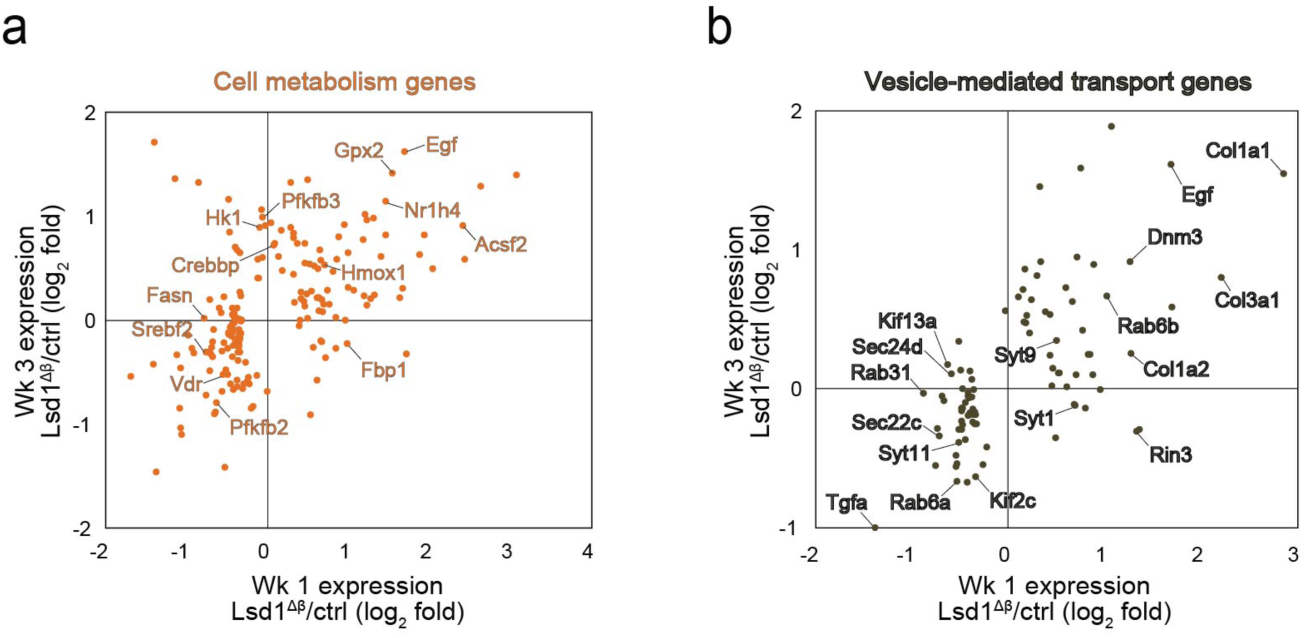
Lsd1 regulates metabolic and vesicle transport genes in β-cells. Related to Figure 4. **(a** and **b)** Log_2_ fold changes in mRNA levels between Lsd1^Δβ^ and control (ctrl) islets at one and three weeks following TM treatment. Genes differentially expressed (*P* < 0.01 by Cuffdiff) in Lsd1^Δβ^ islets compared to control islets at either one or three weeks following TM treatment and annotated to the shown functional categories (from Fig. 4e) are plotted.

**Extended Data Figure 7.**
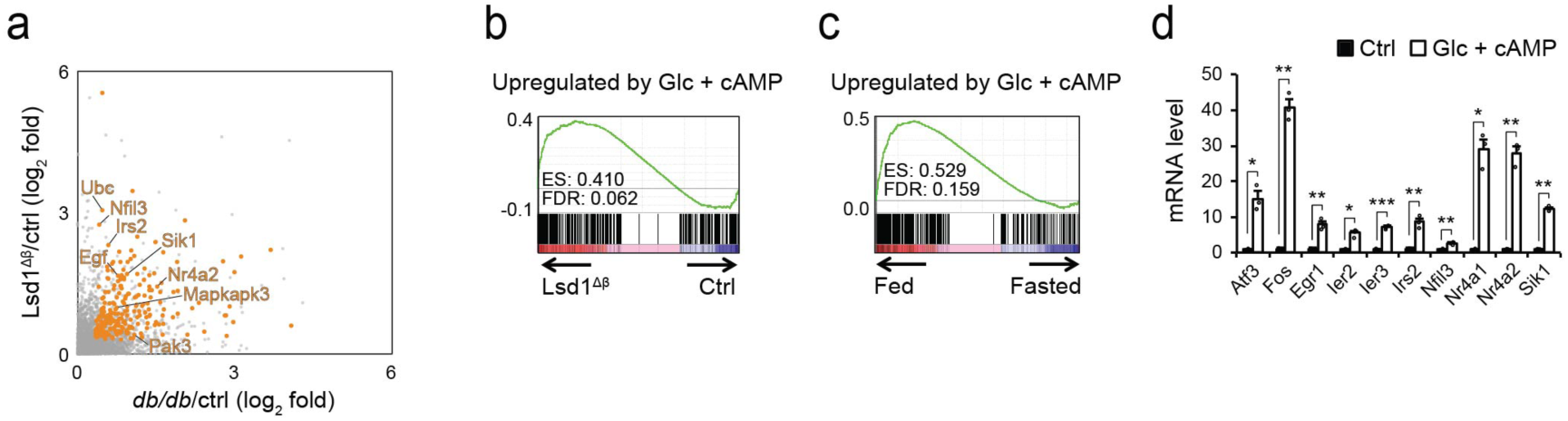
*Lsd1* inactivation upregulates a feeding-induced gene expression signature downstream of cAMP. Related to Figure 4. **(a)** Log_2_ fold changes in mRNA levels in *db/db* compared to control (ctrl, *db*/+) islets and in Lsd1^Δβ^ and control (TM-treated *Pdx1-CreER* mice) islets one week following TM treatment. Genes upregulated in both conditions (*P* < 0.01 by Cuffdiff) are highlighted in orange. For clarity, only the upper right quadrant is shown. **(b** and **c)** Gene set enrichment analysis of genes upregulated by glucose (Glc) and cAMP co-treatment in Min6 insulinoma cells^21^ against mRNA-seq data from Lsd1^Δβ^ and control islets one week following TM treatment (b) or against islet mRNA-seq data from fed and fasted mice (c). **(d)** qPCR of select nutrient-dependent signalling pathway genes in primary mouse islets treated with 11 mM glucose and 0.5 mM cpt-cAMP for 1 hour. *n* = 3 pools of 50 islets each per group. Data are shown as mean ± S.E.M. \**P* < 0.05, \*\**P* < 0.01, \*\*\**P* < 0.001 by unpaired two-tailed t-test with Welch’s correction for unequal variance as necessary. Source data for all quantifications and exact *P*-values for all indicated statistical tests are provided online.

**Extended Data Figure 8.**
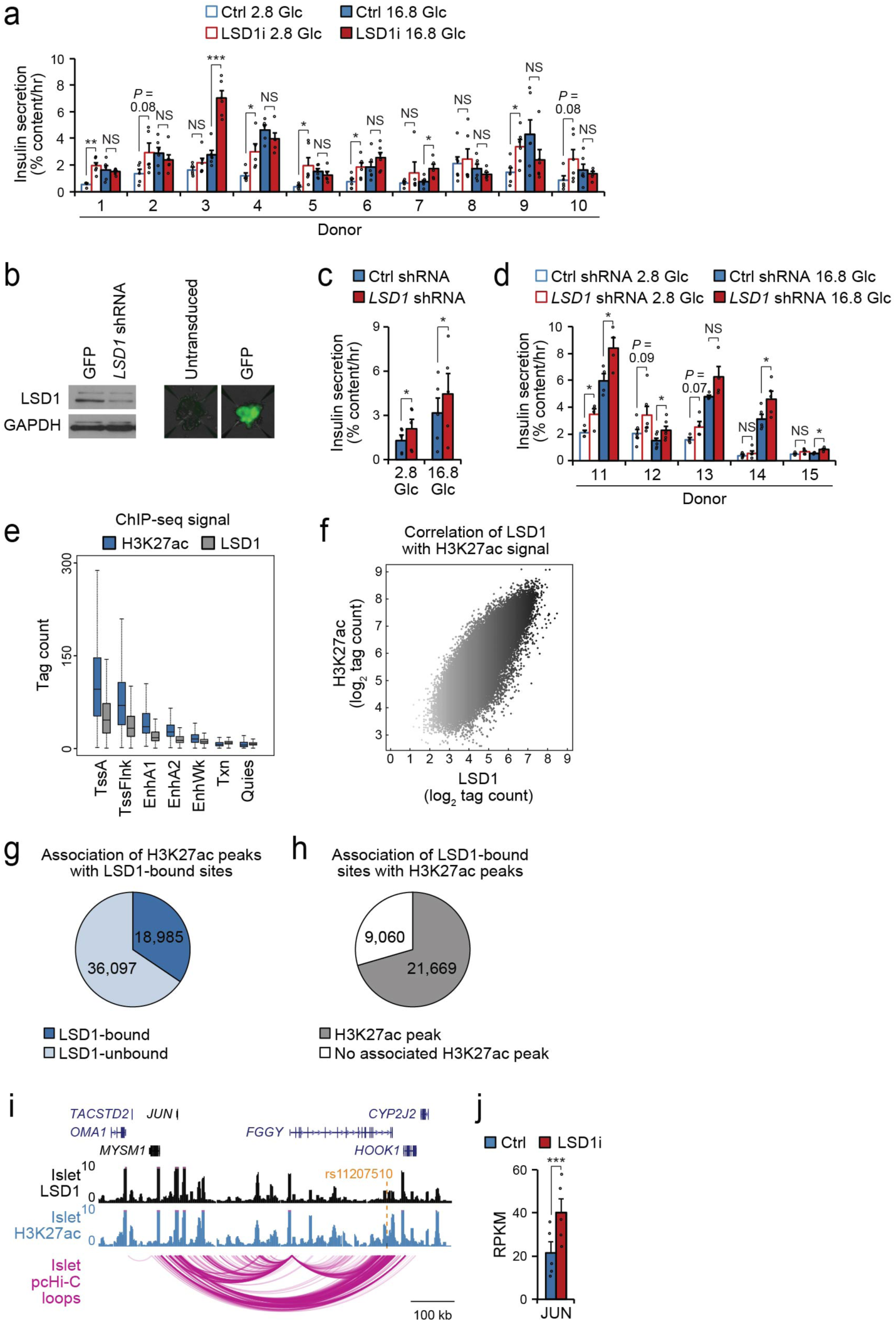
LSD1 associates with active chromatin in human islets. Related to Figure 5. **(a)** Static insulin secretion assays for each human islet preparation stimulated with the indicated glucose (Glc) concentrations (in mM) following 24-hour treatment with the LSD1 inhibitor (LSD1i) TCP or vehicle control (ctrl). Donor 1 2.8 mM Glc + vehicle and 16.8 mM Glc + vehicle, Donor 2 2.8 mM Glc + vehicle, Donor 3 2.8 mM Glc + vehicle, Donor 4 all groups, donor 5 16.8 mM Glc + LSD1i, donor 7 2.8 mM Glc + LSD1i and 16.8 mM Glc + LSD1i, donor 10 2.8 mM Glc + vehicle: *n* = 5 pools of 10 islets each, all other groups: *n* = 6 pools of islets. Data are shown as mean ± S.E.M. \**P* < 0.05, \*\**P* < 0.01, \*\*\**P* < 0.001 by unpaired two-tailed t-test with Welch’s correction for unequal variance as necessary. NS, not significant. **(b)** Western blot for the indicated proteins (left) and GFP fluorescence overlaid with the transmitted light channel (right) in dissociated and reaggregated human islets transduced with *LSD1* shRNA lentivirus or GFP-expressing control lentivirus. **(c** and **d)** Static insulin secretion assays for sorted and reaggregated human β-cells stimulated with the indicated glucose (Glc) concentrations (in mM) following transduction with *LSD1* shRNA or non-targeting (ctrl) shRNA lentiviruses shown in aggregate (c, *n* = 5 donors) or as individual donors (d). Donors 11, 13, and 15: *n* = 4 pools of 5,000 reaggregated β-cells each, donor 12: *n* = 6 pools of reaggregated β-cells, donor 14: *n* = 5 pools of reaggregated β-cells. Data are shown as mean ± S.E.M. \**P* < 0.05 by paired (c) or unpaired (d) two-tailed t-test with Welch’s correction for unequal variance as necessary. NS, not significant. **(e)** H3K27ac and LSD1 ChIP-seq signals at the indicated ChromHMM states in human islets. Data from independent ChIP-seq experiments from four donors (H3K27ac) or 2 donors (LSD1) were merged for analysis. Boxplot whiskers span data points within the interquartile range x 1.5. **(f)** Scatterplot of ChIP-seq signals for H3K27ac and LSD1 in human islets. **(g** and **h)** Numbers of H3K27ac peaks associated with an LSD1 peak ± 1 kb (g) and vice versa (h). **(i)** LSD1 and H3K27ac ChIP-seq and promoter capture Hi-C (pcHi-C)^23^ genome browser tracks showing interaction between the LSD1-bound site containing T2D-associated variant rs11207510 (highlighted in orange) and the *JUN* promoter in human islets. **(j)** Bar graph of *JUN* mRNA level in human islets treated with LSD1i as in (Fig. 5a). Data shown as mean ± S.E.M. \*\*\**P* < 0.001 by Cuffdiff. Source data for all quantifications and exact *P*-values for all indicated statistical tests are provided online.

